# Histone marks are drivers of the splicing changes necessary for an epithelial-to-mesenchymal transition

**DOI:** 10.1101/2021.05.04.442453

**Authors:** A. Segelle, Y. Núñez-Álvarez, A. J. Oldfield, K. M. Webb, P. Voigt, R. F. Luco

## Abstract

Cell differentiation and reprogramming depend on coordinated changes in specific alternative splicing events. How these cell type-specific splicing patterns are dynamically modified in response to a stimulus remains elusive. Taking advantage of the epithelial-to-mesenchymal transition (EMT), a reversible cell reprogramming intimately involved in cancer cell invasiveness and metastasis, we found a strong correlation between changes in the alternative splicing of key exons for EMT, such as at the *Fgfr2* and *Cnntd1* loci, and changes in the enrichment levels of specific histone modifications, namely H3K27ac and H3K27me3. Localised CRISPR epigenome editing of these exon-specific histone marks was sufficient to induce changes in splicing capable of recapitulating important aspects of EMT, such as a motile and invasive cell phenotype. Whereas, impairment of the changes in H3K27 marks observed during EMT, using histone deacetylase inhibitors, repressed inclusion of the mesenchymal isoform despite an EMT induction, supporting a driving effect for H3K27 modifications in establishing the new cell type-specific splicing patterns necessary for EMT cell reprogramming. Finally, H3K27 marks were shown to impact splicing by modulating recruitment of the splicing factor PTB to its RNA binding sites, suggesting a direct link between chromatin modifications and the splicing machinery. Taken together, these results prove the causal role of H3K27 marks in driving the dynamic splicing changes necessary for induction of important aspects of EMT. They also prove that chromatin-mediated splicing changes are sufficient to impact the cell’s phenotype, which expands the cell’s toolkit to adapt and respond to diverse stimuli, such as EMT induction.

## Introduction

During cell reprogramming, such as in the epithelial-to-mesenchymal transition (EMT), cell type-specific transcriptional and splicing programs are tightly regulated to gain new phenotypic traits^1, 2^. Alternative splicing depends on the combinatorial recruitment of specific splicing factors to their corresponding RNA binding sites, which impacts the final splicing outcome^3^. It has long been known that nucleosome positioning and chromatin modifications can modulate this recruitment by impacting RNA polymerase II elongation rate^4^. More recently, results from our laboratory and others found an alternative mechanism of chromatin-mediated splicing regulation. Histone marks were shown to modulate the recruitment of specific splicing factors to weaker RNA-binding sites via protein-protein interactions with chromatin-binding proteins that act as adaptors between the chromatin and the splicing machinery^5, 6^. In this recruitment model, a histone mark, such as H3K36me3, can impact the recruitment of more than one splicing regulator, such as PTB, SRSF1 or EFTUD2 (U5 snRNP), via different chromatin adaptor proteins, like MRG15, PSIP1 or BS69, respectively^7–9^. On the other hand, a splicing factor, such as U2 snRNP, can be recruited by more than one histone mark/chromatin adaptor complex, like H3K4me3/CHD1^10^ and acetyl H3/Gcn5^11^, adding an extra regulatory layer to the alternative splicing reaction for increased specificity and fine-tuning. At a more global level, recent epigenomic analyses have uncovered a coordinating role for histone modifications in regulating the alternative splicing of specific subsets of genes with common regulatory functions^12^. For instance, in acute myeloid leukaemia cell lines, a subset of alternatively spliced exons intimately involved in cell proliferation and transformation were shown to be dependent on local enrichment of H3K79me2^13^. Whereas during stem cell differentiation, exons involved in cell cycle progression and DNA damage response were specifically marked by H3K36me3 and H3K27ac^14^. However, most of this evidence is just correlative, or based on genome-wide alteration of the histone mark of interest via drug-based inhibition and/or overexpression/repression of the chromatin regulator involved, which limits the capacity to properly assess the direct role of a localized histone mark in driving cell type-specific splicing programs important for cell identity. Neither do we know how dynamic changes in splicing are rapidly regulated to establish a novel cell type-specific splicing program in response to a specific stimulus, such as in EMT reprogramming.

Based on our previous results on the alternatively spliced model gene Fibroblast Growth Factor Receptor 2 (*Fgfr2*)^8, 15^, we are now addressing the dynamic role of epigenetic marks in driving the changes in splicing necessary to impact cell biology. To do so, we took advantage of the well-established Epithelial-to-Mesenchymal Transition (EMT), in which changes in the alternative splicing of specific genes, such as *Fgfr2* or *Ctnnd1*, are sufficient to induce cell reprogramming^2, 16, 17^. Temporal correlations during EMT identified H3K27me3 and H3K27ac as the two histone marks for which local changes at alternatively spliced exons essential for EMT preceded the changes in splicing. CRISPR/dCas9 epigenome editing of these H3K27 marks, precisely at the alternatively spliced exon of interest, was sufficient to induce inclusion of the mesenchymal-specific splicing isoform in human epithelial cells. This was done by regulating the recruitment of the splicing regulator PTB to the pre-mRNA, supporting a direct effect of H3K27 marks on the splicing machinery. Additionally, inhibition of H3K27ac changes during EMT impaired inclusion of the mesenchymal-specific splicing event regardless of EMT induction, proving the dominant effect of these histone marks in establishing the new EMT-specific splicing program. Finally, epigenetically induced changes in splicing were sufficient to recapitulate important aspects of the EMT, which supports a major role for histone marks in inducing phenotypically driving splicing changes. These findings uncover a new regulatory layer through which dynamic changes in splicing are regulated by chromatin-dependent mechanisms in response to a specific stimulus, such as in cellular reprogramming.

## Results

### Specific histone modifications correlate in time with dynamic changes in splicing during EMT

The epithelial-to-mesenchymal transition is a cell reprogramming process involved in early development, wound healing, and tumour invasion in metastasis^1, 2^. Human epithelial MCF10a cells stably expressing the EMT inducer SNAIL1 fused to the oestrogen receptor (MCF10a-Snail-ER) can be reprogrammed into mesenchymal-like cells in less than a week by addition of the ER ligand tamoxifen (Figure 1A)^1^. The first changes in splicing of classical EMT genes, such as *Fgfr2, Ctnnd1, Slk and Scrib*, were observed as early as 12h after induction (T0.5) (Figure 1C,H and S1F,N,Q). Moreover, all changes in splicing could be reversed, through a mesenchymal-to-epithelial transition (MET), by removing tamoxifen from the culture medium for three weeks, highlighting the dynamic nature of this cellular system (Figure 1C,H and S1F). When comparing changes in alternative splicing of key EMT genes (*Ctnnd1, Enah, Fgfr2, Slk, Scrib* and *Tcf7l2*, Figure 1A) to changes in histone modifications levels previously shown to mark alternatively spliced genes^8, 15^, we found that changes in H3K27me3, H3K27ac and H3K4me1 strongly correlated in time with splicing changes in 5 out of 6 genes studied (Figure 1B-K and S1I-R). However, H3K4me1 correlated rather at late phases of EMT (Figure 1F,K, S1I,J,L,M), and H3K36me3 rarely showed a correlation with changes in splicing (Figure S1D,H and data not shown). These epigenetic changes were highly localised, occurring precisely over the alternatively spliced exon (Figure 1D-F,I-K and S1). H3K27me3 and H3K27ac levels were anti-correlated in 3 out of the 5 genes analysed, while H3K4me1 changed rather in the same direction as H3K27ac, suggesting distinct combinatorial effects in splicing regulation (Figure 1D-F,I-K and S1I,J). Furthermore, with the exception of the mutually exclusive exons in *Fgfr2*, H3K27me3 levels positively correlated with inclusion of all the alternatively spliced exons tested, which points to a regulatory role in coordinating a specific splicing program during EMT (Figure 1D and S1I,J,L,M,P). Of note, these changes in exon-specific histone marks did not correlate with changes in gene expression nor nucleosome positioning during EMT (Figure S1A,C,E,G), suggesting a splicing-specific effect. Finally, the observed changes in chromatin modifications were not only as dynamic as the changes in splicing, but also reversible upon MET, implying epigenetic plasticity (Figure 1, MET panel).

**Figure 1:**
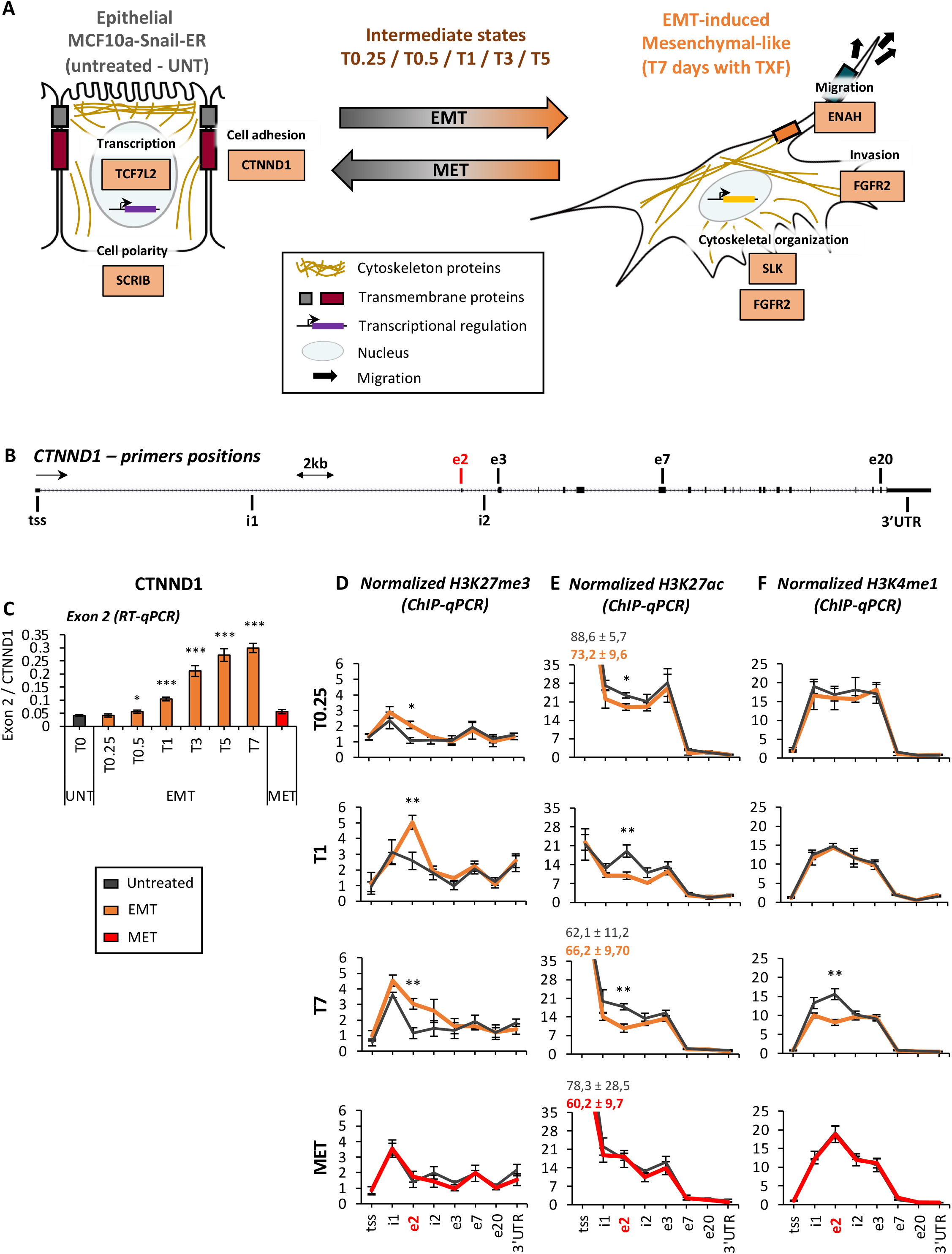
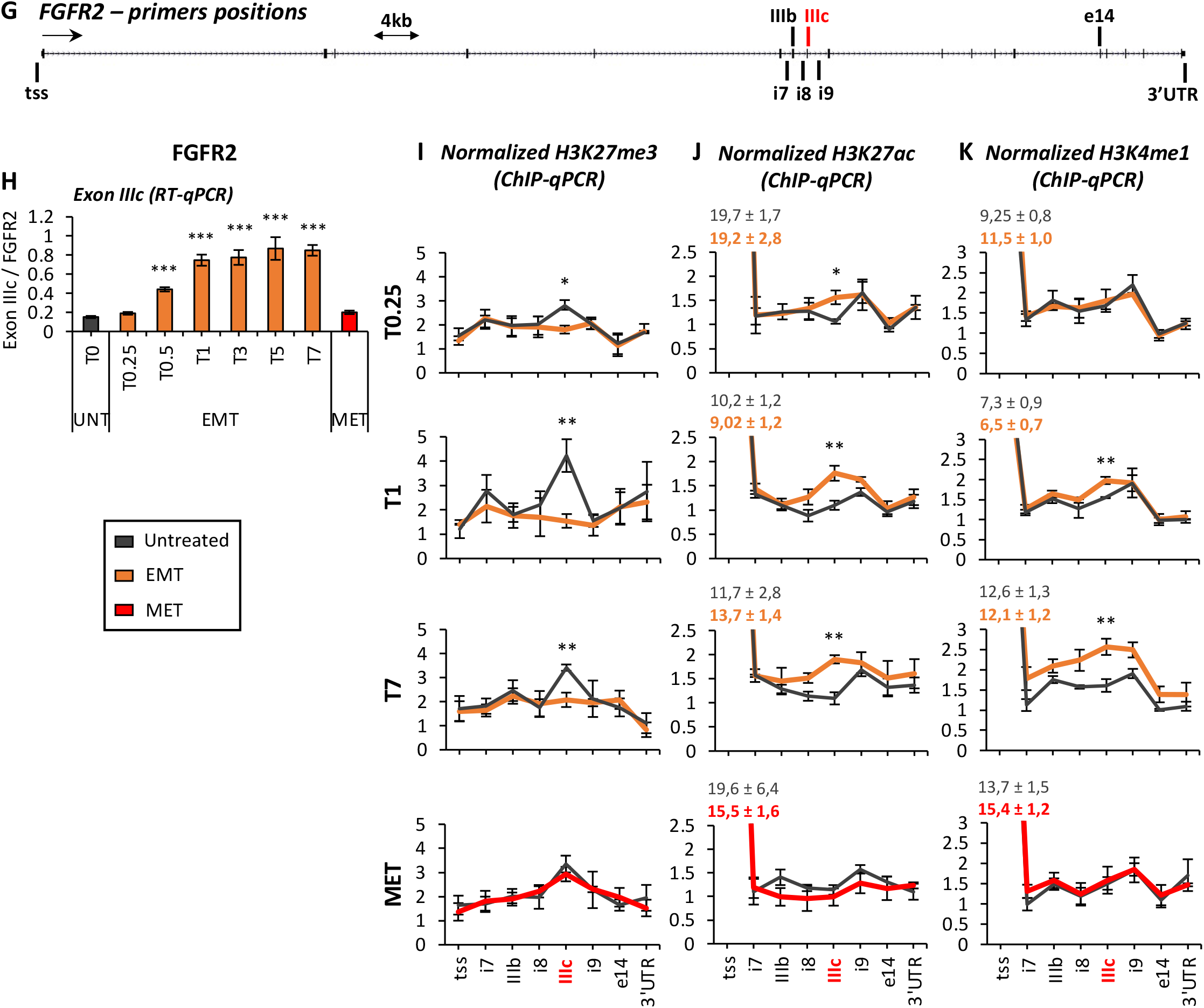
Specific histone modifications correlate in time with dynamic changes in splicing during EMT. **(A)** Schematic representation of the epithelial-to-mesenchymal transition (EMT) and reverse MET. The function of the alternatively spliced genes most relevant for EMT transition is shown. Normal human epithelial MCF10a-Snail-ER are totally reprogrammed into mesenchymal-like cells in 7 days (T7). First changes in EMT markers are observed 6h (T0.25) after treatment with tamoxifen (TXF). Until the EMT induction is complete, there are several intermediate states in which heterogenous populations of cells coexist. (**B,G**) Representation of CTNND1 and FGFR2 gene loci in which the position of the primers used for ChIP-qPCR experiments is indicated. Highlighted in red are the alternatively spliced exons regulated during EMT (**C, H**) Inclusion levels of CTNND1 exon 2 and FGFR2 exon IIIc relative to total expression levels of CTNND1 and FGFR2, respectively, in MCF10a-Snail-ER cells at different time points during induction of the EMT (0 to 7 days in presence of tamoxifen, orange) and reversible MET (21 days after removal of the tamoxifen at T7, red). RT-qPCR results are shown as the mean +/- SEM of n=4 biological replicates. (**D-F** and **I-K**) Enrichment levels of H3K27me3 (D, I), H3K27ac (E, J) and H3K4me1 (F, K) along CTNND1 or FGFR2 locus, focusing into the alternatively spliced exons of interest (CTNND1.e2 and FGFR2.IIIc) and flanking intronic and exonic control regions, in tamoxifen-induced MCF10a-Snail-ER cells treated for 6h (T0.25), 24h (T1) or 7 days (T7) with tamoxifen and MET reversed cells in which the tamoxifen was eliminated for 21 days (MET). Chromatin immunoprecipitation results are shown as the mean +/- SEM in n=4 biological replicates. The percentage of input was normalized by two control regions across the different conditions. *P <0.05, **P <0.01, ***P <0.001 in two-tail paired Student’s t-test respect untreated cells (grey).

In conclusion, we have found a localised enrichment of specific histone marks, H3K27me3, H3K27ac and H3K4me1, whose changes correlate in time with the highly dynamic splicing changes observed during the reprogramming of an epithelial cell into a mesenchymal one during EMT, which points to a potential functional link.

### Localised changes in H3K27me3 and H3K27ac are sufficient to induce exon-specific changes in alternative splicing

In contrast with H3K4me1 late changes during EMT, H3K27me3 and H3K27ac changes were evident prior to the detection of splicing changes in *Ctnnd1* and *Fgfr2* genes, at 6h post-induction (Figure 1, T0.25 panel), suggesting a causative effect of these marks on alternative splicing. To directly test this hypothesis, we adapted the CRISPR/dCas9 system^18^ to edit the epigenome specifically at differentially marked, alternatively spliced exons. Catalytic domains of well-known H3K27 modifiers were fused to a DNA targeting-competent, but nuclease-dead, mutant dCas9 to induce site-specific changes in H3K27 methyl or acetyl levels. Using this system, EZH2 H3K27 methyltransferase^19^, UTX1 demethylase^20^, p300 acetyltransferase^18^ and Sid4x deacetylase^21^ were targeted to CTNND1 exon 2 or FGFR2 exon IIIc in untreated epithelial MCF10a-Snail-ER cells (Figure 2A,G). To verify the exon specificity of the system, alternatively spliced exons present in the same gene, but not differentially enriched for these histone marks during EMT, namely CTNND1 exon 20 and FGFR2 exon IIIb, were also targeted using the same dCas9 modifiers (Figure S2C,L). As expected, dCas9-p300, but not its catalytic mutant dCas9-p300*, increased H3K27ac levels specifically at the targeted exons in both genes (Figure 2B,H and S2D,H,M,Q). On the other hand, dCas9-Sid4x slightly reduced H3K27ac levels just at CTNND1 exon 2 and dCas9-UTX1 reduced H3K27me3 levels mostly at exon IIIc, which are the exons enriched in these marks in epithelial cells (Figure 2B,C,H,I). dCas9-EZH2, though, had only a minor effect on H3K27me3 levels. To improve H3K27me3 editing, we tested vSET, a viral SET domain protein that specifically methylates H3K27 without requiring Polycomb Repressive Complex 2 subunits for activity (Figure S2A-B)^22^. As expected, a dimeric vSET construct fused to dCas9 (dCas9-vSETx2), but not its catalytic mutant dCas9-vSETx2*, strongly increased H3K27me3 levels precisely at the targeted exons (Figure 2C,I and S2E,I,N,R). Finally, H3K4me1, H3K9ac and H3K9me2 were not affected by H3K27 epigenome editing, confirming the specificity of the system (Figure S2U,V). Interestingly, the increase in H3K27ac levels, mediated by dCas9-p300, also resulted in reduced H3K27me3 levels, while dCas9-UTX1-mediated H3K27 demethylation increased H3K27ac levels and dCas9-vSETx2 reduced H3K27ac (Figure 2B,C,H,I and S2D). These findings confirm the anti-correlative nature of these marks and establish the capacity of the CRISPR-dCas9 system to generate the chromatin signatures observed during EMT.

**Figure 2:**
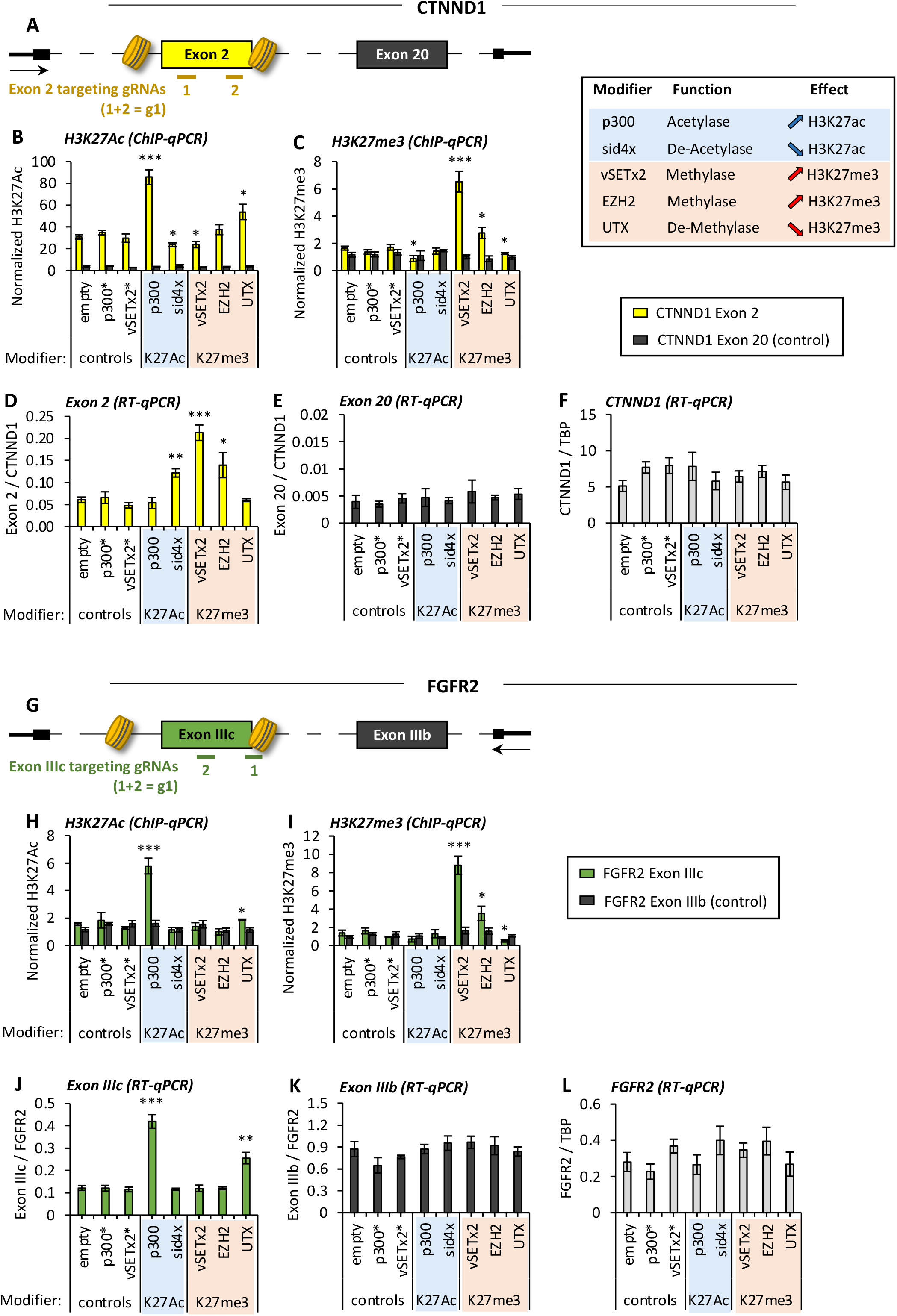
Exon-specific epigenome editing of H3K27 marks is sufficient to induce a change in splicing. (**A**) Schematic representation of CTNND1 gene locus and alternatively spliced exon 2 (yellow) and exon 20 (grey). The position of the gRNAs used to exon-specifically target the different dCas9-fused proteins are represented in colour-coded lines. Nucleosome positioning, according to MNase-qPCR assay (data not shown), is shown in exon 2. (**B,C**) Enrichment levels of H3K27ac (B) and H3K27me3 (C) at CTNND1 exon 2 (yellow) and control exon 20 (grey) in MCF10a-Snail-ER cells upon infection of dCas9 fused to the catalytic domain of an H3K27 epigenetic modifier (see summary table on the right) in the presence of exon-specific gRNAs targeting exon 2, by quantitative chromatin immunoprecipitation (mean +/- SEM, n=4). The percentage of input was normalized by two control regions across the different conditions. Mutated p300* and vSETx2* were used as negative controls together with empty dCas9. (**D-F**) Expression levels of CTNND1 exon 2 (D), exon 20 (E), and total CTNND1 (F) relative to total TBP and CTNND1 levels, respectively, in MCF10a-Snail-ER cells upon infection with dCas9 H3K27 epigenome editors and exon 2-specific gRNAs, determined by quantitative RT-qPCR (mean +/- SEM, n=4). (**G**) Schematic representation, as in (A), of FGFR2 gene, gRNAs position at the targeted exon IIIc (green) and nucleosome positioning at exon IIIc (data not shown). (**H,I**) H3K27ac and H3K27me3 enrichment levels at the gRNA-targeted exon IIIc and control IIIb in MCF10a-Snail-ER cells infected with the dCas9 H3K27 modifiers by ChIP as described in (B,C) (mean +/- SEM, n=4). (**J-L**) Expression levels of exon IIIc, control IIIb and total FGFR2, relative to total TBP and FGFR2 expression levels, respectively, by quantitative RT-qPCR as described in D-F (mean +/- SEM, n=4). *P <0.05, **P <0.01, ***P <0.001 in two-tail paired Student’s t-test respect empty dCas9 plasmid (empty).

As predicted from the changes in histone modifications observed during EMT (Figure 1D,E), only dCas9-EZH2/vSETx2-mediated increase in H3K27me3 levels, or decrease in H3K27ac using dCas9-Sid4x, affected CTNND1 splicing, resulting in a ∼3.5x-fold increase in the inclusion of the mesenchymal-specific exon 2 (Figure 2D). In contrast, consistent with the exon-specific H3K27 signature observed in FGFR2 exon IIIc in EMT cells (Figure 1I,J), exon IIIc splicing was induced by an increase in H3K27ac (dCas9-p300) and by a decrease in H3K27me3 (dCas9-UTX1) levels (Figure 2J). These results proved the driving effect of these histone marks in inducing specific splicing changes. Importantly, H3K27 epigenome editing did neither affect the total expression levels of these genes, nor splicing of other exons, such as CTNND1 exon 20 or FGFR2 exon IIIb, supporting an exon-specific splicing effect (Figure 2E,F,K,L). Furthermore, the use of catalytically dead mutants, such as dCas9-vSETx2* and dCas9-p300*, did not impact exon inclusion levels either, which validated an epigenetic-dependent splicing effect (Figure 2D,J). Finally, targeting CTNND1 exon 2 or FGFR2 exon IIIc with a second set of gRNAs (g2) also consistently induced inclusion of the mesenchymal-specific isoforms, when the corresponding dCas9 modifier was used, confirming the robustness of the results and ruling out potential off-target effects (Figure S2C-G, L-P). It is important to note that epigenome editing of alternatively spliced exons not differentially marked by H3K27 modifications during EMT, such as CTNND1 exon 20, FGFR2 exon IIIb or ENAH exon 11 had no impact on their splicing (Figure S2H-K, Q-T and not shown), suggesting a context-specific regulatory effect at exons marked by H3K27 modifications.

Taken together, local changes in specific histone modifications at epigenetically-marked exons are sufficient to trigger the dynamic changes in alternative splicing observed during EMT, which supports a causal role for chromatin marks on inducing cell type-specific splicing changes. We next sought to study the importance of these dynamic epigenetic changes in splicing reprogramming during EMT induction.

### Dynamic changes in H3K27ac and H3K27me3 are necessary to induce a change in splicing during EMT

To test whether the changes in H3K27 marks are necessary to drive the changes in splicing observed during EMT, MCF10a-Snail-ER cells were treated with a histone pan-deacetylase inhibitor (HDACi) during EMT induction. As expected, both Trichostatin A (TSA) and the less toxic Panobinostat (Pano)^23^ maintained H3K27ac levels significantly higher than in control cells (DMSO) at the exons of interest during EMT induction (Figure S3A and not shown). Despite a successful EMT (Figure S3B and not shown), exons in which there is a depletion in H3K27ac levels and/or increase in H3K27me3 during EMT, which are CTNND1 exon 2, SCRIB exon 16 and to a less extent SLK exon 13, did not shift to the expected mesenchymal-specific splicing isoform (Figure 3A-C). In contrast, exons with no changes in H3K27 marks during EMT (ENAH, CLSTN1, PLOD2) were not impacted, or just impacted by one of the HDAC inhibitors, supporting a specific effect on exons sensitive to H3K27ac (Figure 3D-F). To reduce pleiotropic indirect effects from the inhibitors (Figure S3C) and narrow down the HDACs necessary for dynamic changes in splicing, we specifically knocked-down catalytically active HDACs expressed in MCF10a-Snail-ER cells. Neither HDAC1, HDAC2, HDAC3 nor HDAC8 knockdown had an effect on *Ctnnd1*, *Scrib* nor *Slk* splicing (data not shown). However, due to known redundancy between HDAC1 and HDAC2^24^, we performed a double knock-downed of both deacetylases with lentiviral shRNAs. Even though HDAC1 could not be repressed to more than 60% (Figure S3D), co-repression of the two HDACs prior to EMT induction recapitulated HDACi results, without impacting expression of any of the splicing regulators of relevance for EMT, confirming the driving role of H3K27 marks in inducing specific changes in alternative splicing (Figure 3G-L and S3F).

**Figure 3:**
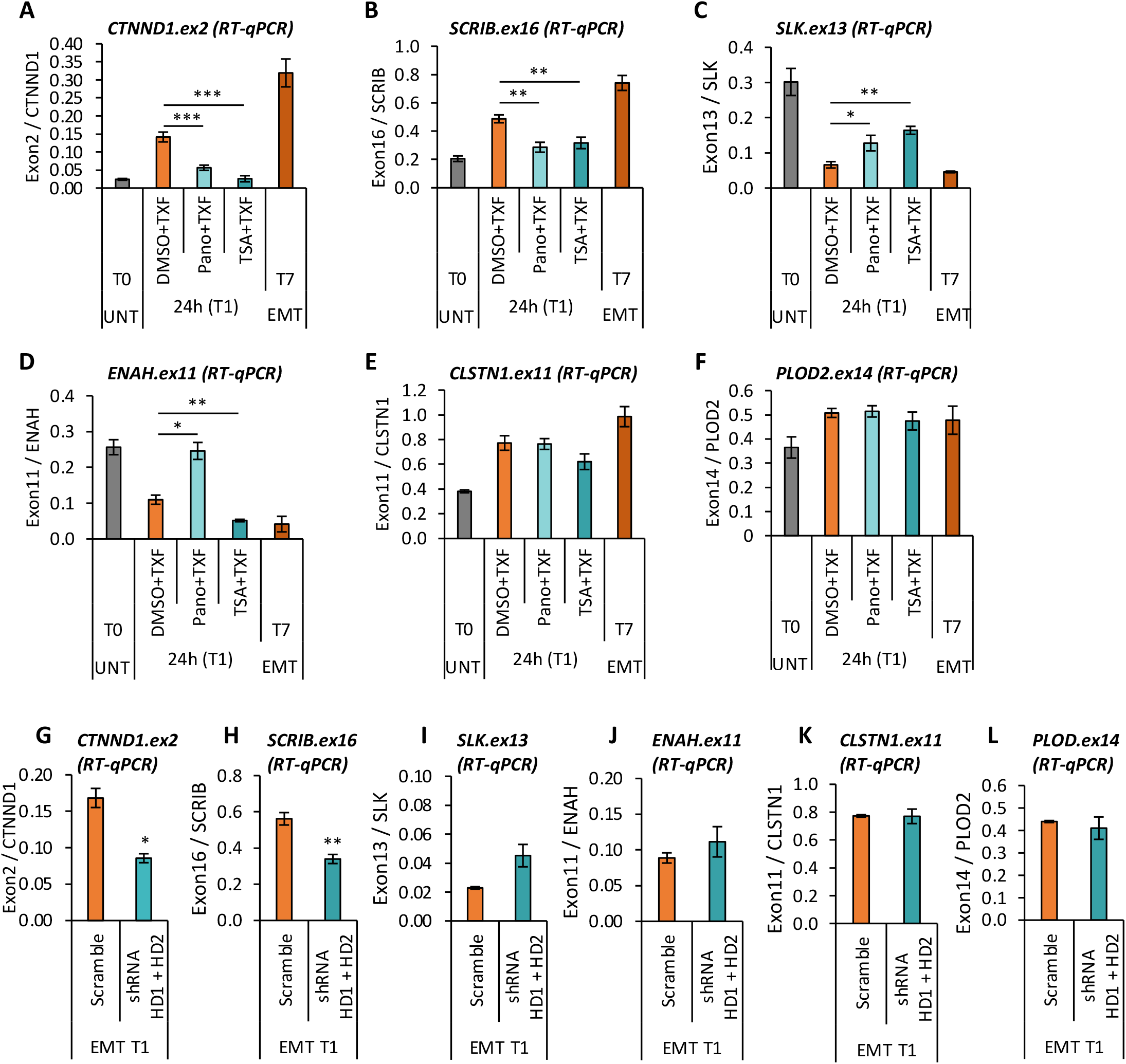
HDAC inhibition during EMT induction prevents the shift to the mesenchymal-specific isoform at alternatively spliced genes differentially marked by H3K27ac/me3. (**A-F**) Inclusion levels of H3K27-marked exons (CTNND1, SCRIB, SLK) relative to total expression levels of the corresponding gene in MCF10a-Snail-ER cells treated for 24h with tamoxifen for EMT induction (T1, orange) and 10nM of Pano (Panobinostat, light cyan), 3µg/mL of TSA (Trichostatin A, cyan) or control vehicle (DMSO, orange) for inhibition of the changes in H3K27ac observed during EMT. ENAH, CLSTN1 and PLOD were used as control genes. Untreated (T0, grey) and fully induced (T7, orange) EMT cells are shown as control references. RT-qPCR results are shown as the mean +/- SEM of n=3 biological replicates. (**G-L**) Inclusion levels of the same exons as before relative to total expression levels of the corresponding gene in MCF10a-Snail-ER cells at day 1 of EMT induction upon double knock-down of HDAC1 (HD1) and HDAC2 (HD2). Non-targeting shRNA (scramble) is used as a control. RT-qPCR results are shown as the mean +/- SEM of n=3 biological replicates. *P <0.05, **P <0.01, ***P <0.001 in two-tail paired Student’s t-test respect T1 DMSO for (A-F) and T1 Scramble for (G-M).

In conclusion, changes in H3K27ac/me3 levels are sufficient and necessary to induce the dynamic changes in splicing observed during EMT. We next sought to understand how H3K27 marks regulate splicing using *Ctnnd1* as a model gene.

### H3K27 marks do not regulate splicing by modulating RNA polymerase II elongation rate

Chromatin has long been proposed to impact splicing by modulating RNA polymerase II elongation rate, which alters the kinetics of splicing factor recruitment to competing alternative splice sites in nascent transcripts^5, 6^. As H3K27me3 is known to mediate chromatin compaction, which can slow down transcription kinetics, and H3K27ac displays opposite effects on chromatin and RNA polymerase II dynamics^4, 19^, we compared RNA polymerase II elongation rates at CTNND1 exon 2 before and after EMT induction in MCF10a-Snail-ER cells. As expected, using the RNA polymerase II inhibitor DRB for synchronous pause / release of transcription in a cell population, we found a delay in transcription of CTNND1 exon 2, but not in the constitutively spliced exon 15, in tamoxifen-induced EMT cells (Figure 4B,E). This delay correlated with enrichment of H3K27me3 and RNA polymerase II at exon 2, which is consistent with a slowdown of RNA polymerase II kinetics (Figure 1D and 4C,F). However, this RNA polymerase II effect was not observed 12h after induction of EMT (T0.5, Figure 4A,D), even though changes in H3K27 marks and exon 2 inclusion were already detected at this time point (Figure 1, T0.25 panels), suggesting that early changes in splicing, dependent on H3K27 marks, are unlikely to be mediated by changes in RNA polymerase II kinetics. Finally, treatment with drugs increasing (TSA) or decreasing (DRB) RNA polymerase II elongation rate did not have an effect on CTNND1 splicing in steady-state epithelial nor EMT-induced mesenchymal-like MCF10a-Snail-ER cells (Figure S4A-B), ruling out an RNA polymerase II-mediated effect.

**Figure 4:**
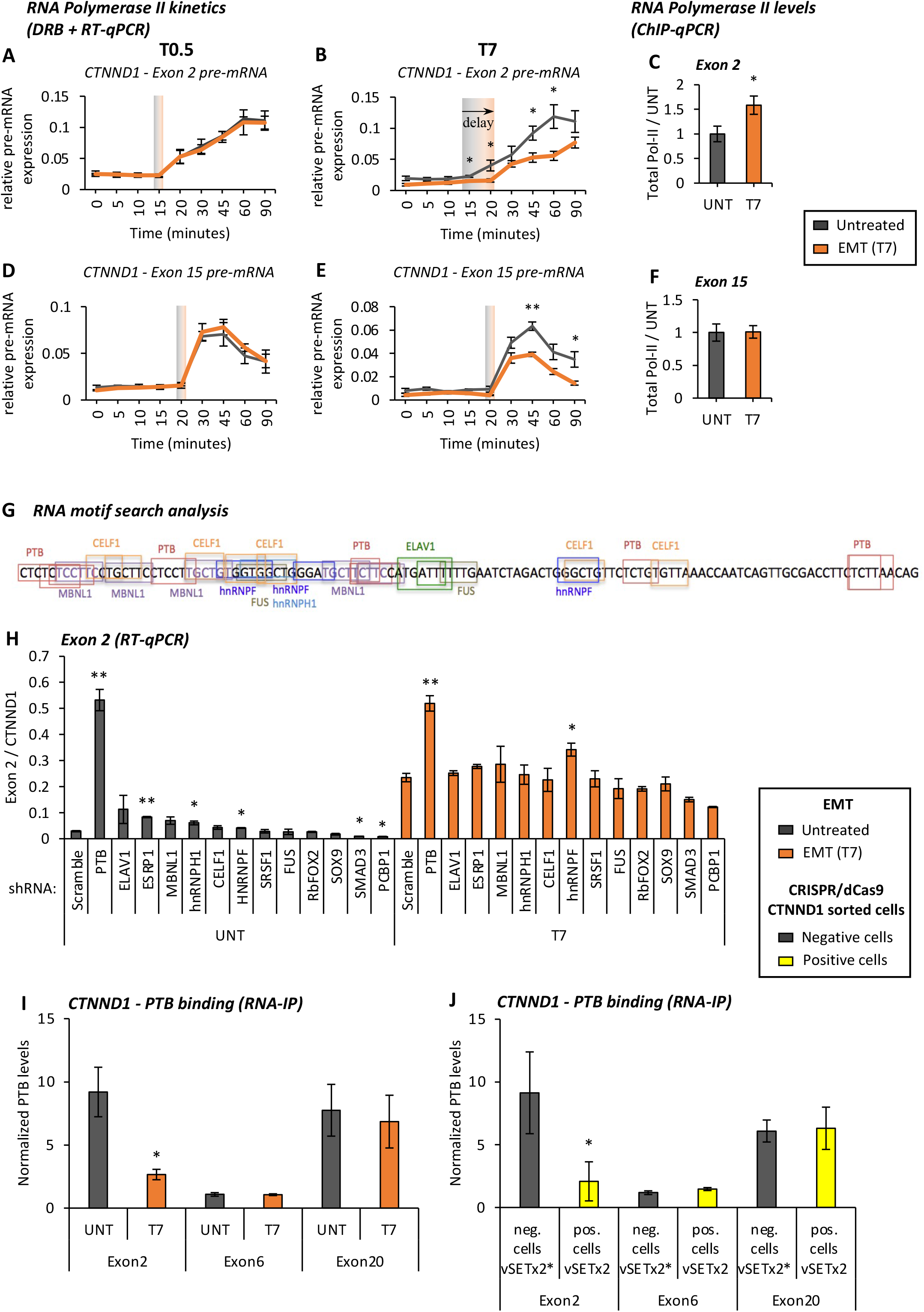
H3K27 marks regulate splicing by modulating the recruitment of specific splicing factors to the pre-mRNA. (**A,B**) Apparition in time of CTNND1 exon 2 pre-mRNA in synchronized untreated (grey) and tamoxifen-induced (orange) MCF10a-Snail-ER cells upon release of the transcriptional inhibitor DRB after 0.5 days (A, T0.5) or 7 days (B, T7) of EMT induction. RT-qPCR results are normalized by tRNA expression levels (mean +/- SEM, n=3). (**C**) Total RNA Polymerase II levels at CTNND1 exon 2 in untreated (grey) and tamoxifen-induced (orange) MCF10a-Snail-ER cells by ChIP-qPCR (mean +/- SEM, n=3). The percentage of input was normalized by two control regions across the different conditions and represented relative to untreated cells (grey). (**D-F**) Same as (A-C) for CTNND1 control exon 15 (mean +/- SEM, n=3). *P <0.05, **P <0.01, ***P <0.001 in two-tail paired Student’s t-test respect untreated cells (grey). (**G**) Predicted RNA-binding motifs along CTNND1 exon 2 pre-mRNA in at least two of the four software used (RBPDB, RBPMAP, SFMAP and Spliceaid, details in methods). (**H**) CTNND1 exon 2 inclusion levels upon knock down, using lentiviral shRNAs, of candidate splicing regulators in untreated (grey) and tamoxifen-induced MCF10-Snail-ER (T7, orange) cells. RT-qPCR results are normalized by total CTNND1 expression levels (mean ± SEM, n=3). (**I**) PTB enrichment levels at CTNND1 exon 2 pre-mRNA in untreated (UNT) and tamoxifen-induced (T7) MCF10a-Snail-ER cells. Constitutively included CTNND1 exon 6 and excluded CTNND1 exon 20 were used as negative and positive controls of PTB binding, respectively. The percentage of input in UV-crosslinking RNA immunoprecipitation was normalized by IgG and CTNND1 exon7 control levels (mean ± SEM, n=5) (**J**) PTB enrichment levels at CTNND1 exon 2 and control exon 6 and exon 20 pre-mRNA in cell-sorted cells expressing (positive) or not (negative) the mesenchymal-specific splicing isoform mCTNND1(ex2) in MCF10a-Snail-ER cells infected with dCas9-vSETx2, or mutant dCas9-vSETx2*, and the exon-specific gRNAs (g1) targeting CTNND1 exon 2. The percentage of input in UV-crosslinking RNA immunoprecipitation was normalized by IgG and CTNND1 exon7 control levels (mean ± SEM, n=6). *P <0.05, **P <0.01 in two-tail paired Student’s t-test respect control cells (scramble shRNA or untreated cells).

We thus conclude that changes in RNA polymerase II elongation rate do not play a role in establishing the CTNND1 mesenchymal isoform during EMT, but may be a consequence of the new cell type-specific splicing pattern that could play a role in its maintenance, as a feed-back mechanism to reinforce the new splice site choice.

### H3K27 marks modulate the recruitment of specific splicing factors to the pre-mRNA

In parallel to the RNA polymerase II kinetic model, we and others have identified a more direct role for histone and DNA marks in regulating the recruitment of splicing regulators to the pre-mRNA^8–10, 25, 26^. To identify the splicing factors involved, we first tested a panel of regulators potentially involved in CTNND1 exon 2 splicing to determine their possible connection with H3K27 marks. shRNA-mediated knockdown of all the RNA binding proteins previously implicated in CTNND1 splicing regulation^27–29^, or identified by motif search analysis, pointed to the splicing factor PTB as the major repressor of CTNND1 exon 2 inclusion (Figure 4G-H and S4C). UV-crosslinking RNA immunoprecipitation assays further revealed differential recruitment of PTB to exon 2 pre-mRNA during EMT, with preferential binding to the H3K27ac-marked exon in untreated epithelial MCF10a-Snail-ER cells, when the exon is excluded (Figure 4I and S4D).

We next assessed the impact of altering H3K27ac/me3 levels, using the dCas9-vSETx2 construct, on PTB recruitment to CTNND1 exon 2 in epithelial MCF10a-Snail-ER cells. As predicted, a local increase in H3K27me3 levels at CTNND1 exon 2, which increases exon inclusion, reduced PTB binding to the exon pre-mRNA. Whereas PTB binding to control regions, such as CTNND1 exon 6 and exon 20, was not affected (Figure 4J and S4E). These findings suggest a direct impact of H3K27 marks on PTB recruitment to CTNND1 exon 2 pre-mRNA.

Of note, all the exons found to be sensitive to H3K27 marks, namely FGFR2 exon IIIb, SCRIB exon 16, SLK exon 13 and TCF7L2 exon 4, were also dependent on PTB levels (Figure S4F-H and ^8^). Even more, PTB knock-down recapitulated the splicing phenotype observed when H3K27ac levels are low and/or H3K27me3 levels are high, supporting a direct effect of H3K27 marks on PTB recruitment (Figure 1, S1 and 4). Genome-wide studies will assess the global impact of H3K27ac and H3K27me3 in coordinating a PTB-dependent splicing program, known to play a major role in EMT and cancer^2^.

We conclude that in the genes studied, dynamic changes in H3K27 marks directly impact alternative splicing by modulating PTB recruitment to the pre-mRNA.

### Chromatin-induced changes in splicing recapitulate the EMT

The physiological impact of chromatin-mediated changes in splicing has long been controversial. Changes in CTNND1 and FGFR2 alternative splicing are important regulators of EMT, and their mesenchymal-specific isoforms have been associated with poor prognosis in several carcinomas, including breast and prostate cancers^2, 30–32^. An increase in CTNND1 exon 2 inclusion levels affects the capacity of the protein to interact with E-cadherins, destabilizing cell-cell interactions and increasing cell motility and invasiveness^17^. Alternatively, an H3K36me3-mediated decrease of FGFR2 exon IIIc mesenchymal isoform, which impacts the ligand specificity of the receptor, was shown to significantly decrease the migratory and invasive phenotype of non-small lung cancer cells, without impacting proliferation or apoptosis^33^. We thus tested whether H3K27-mediated epigenetic induction of the mesenchymal-specific isoforms of CTNND1 or FGFR2 would be sufficient to reproduce migratory EMT-like phenotypes in epithelial MCF10-Snail-ER cells.

dCas9-Sid4x- or dCas9-vSETx2-mediated increase in CTNND1.ex2 mesenchymal isoform and dCas9-p300- or dCas9-UTX1-mediated increase in FGFR2.IIIc mesenchymal isoform significantly decreased the expression of classical epithelial markers, such as E-cadherin and EPCAM, while increasing the expression of the mesenchymal markers ECM-1 and MCAM by ∼2-fold, both at the transcript and protein level (Figure 5A-D). This was specific to the epigenetic editors impacting splicing since none of the other dCas9 modifiers nor dCas9 mutants had an effect (Figure 5A-D and not shown). Furthermore, the H3K27me3-mediated shift in CTNND1 splicing significantly increased the non-directional (wound-healing) and bi-directional (transwell assay) migration capacity of targeted MCF10a-Snail-ER cells (Figure 5E-F), whereas catalytically dead dCas9-p300* and dCas9-vSETx2* had no effect (Figure 5A-F). None of the splicing regulators known to play a role in EMT changed expression levels upon CRISPR epigenome editing (Figure S5A), supporting a direct chromatin-mediated effect on the EMT phenotype. We conclude that highly localised changes in H3K27 marks at alternatively spliced exons important for EMT are sufficient to induce a partial cell reprogramming.

**Figure 5:**
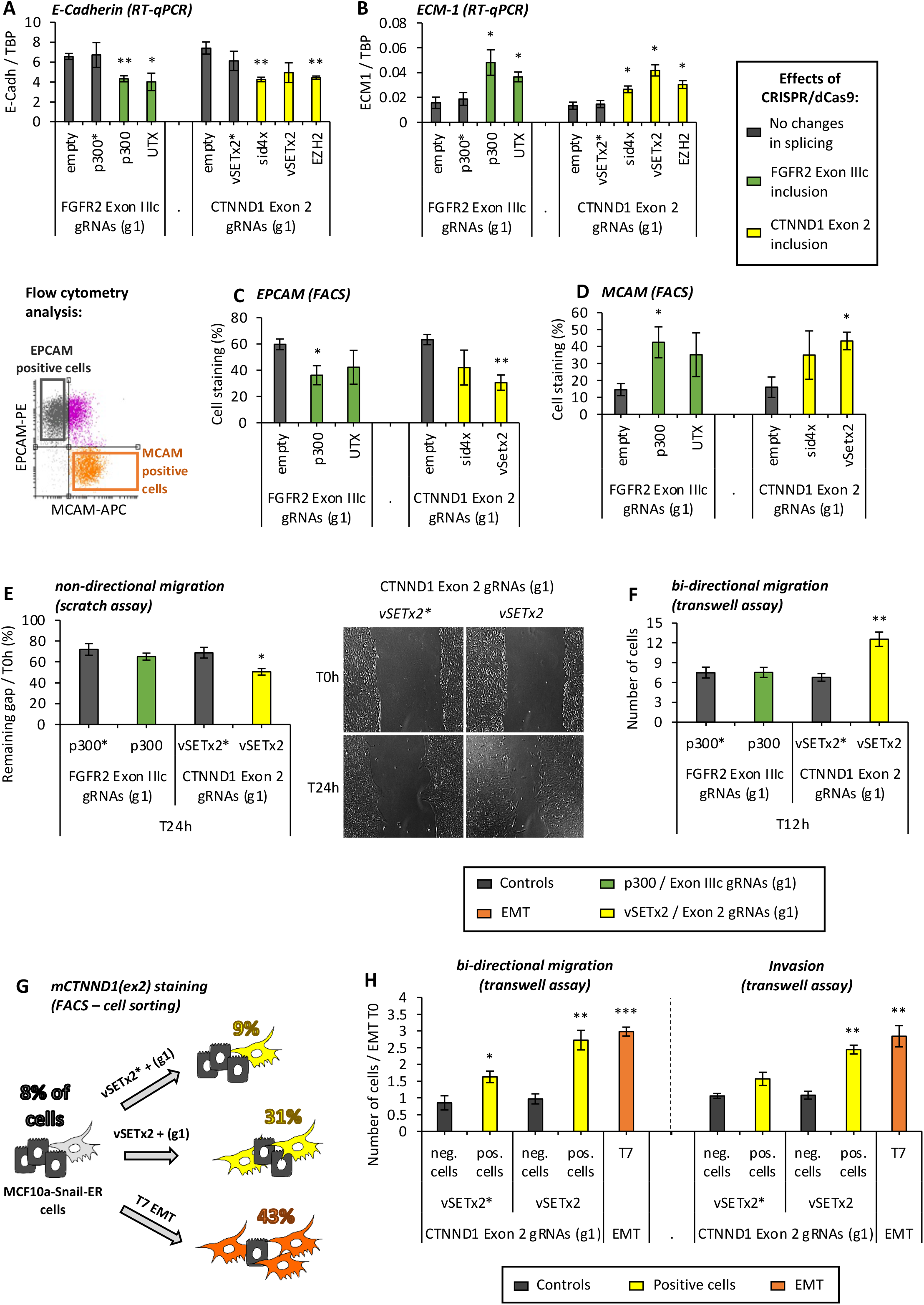
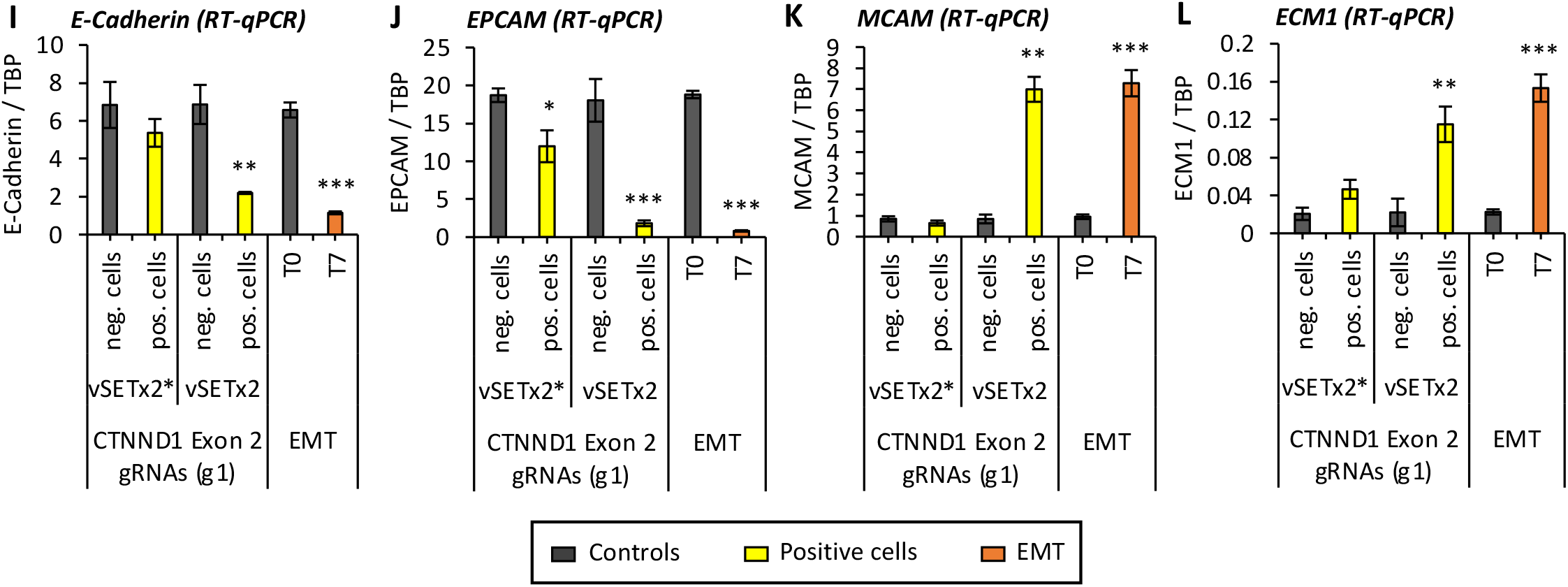
Chromatin-induced changes in splicing recapitulate an EMT. (**A-D**) Expression levels of epithelial (E-Cadherin, EPCAM) and mesenchymal (ECM1, MCAM) markers at the mRNA (A, B - RT-qPCR, mean +/- SEM, n=4) and protein (C,D - Flow cytometry, mean +/- SEM, n=4) levels in MCF10a-Snail-ER cells infected with the dCas9-fused proteins changing splicing and the corresponding exon-specific gRNAs targeting FGFR2 exon IIIc (g1) or CTNND1 exon 2 (g1). mRNA levels are normalized by TBP, and protein levels are quantified above the no primary antibody background signal (summary scheme on the left). (**E-F**) Functional EMT assays to test non-directional (E) and bi-directional migration (F) in MCF10-Snail-ER cells infected with exon 2 or exon IIIc-specific targeting gRNAs and dCas9-fused proteins with their corresponding catalytic mutants. Scratch assays (**E**) were carried out on confluent monolayers of cells for evaluating the % of gap remaining 24h after wound (mean +/- SEM, n=3). Transwell assays (**F**) evaluate the number of cells migrating towards FGF-2 in 12h (mean +/- SEM, n=3). (**G**) MCF10a-Snail-ER cells infected with gRNAs targeting CTNND1 exon 2 and either dCas9-vSETx2 or mutant dCas9-vSETx2* were cell-sorted using a splicing-specific antibody detecting only CTNND1 mesenchymal protein variant (mCTNND1(ex2)) for directional migration and invasion transwell assays. Negative cells not expressing mCTNND1(ex2) and tamoxifen-induced T7 EMT cells were used as control references. The percentage of mCTNND1(ex2) positive cells is shown on the right. (**H**) The number of sorted cells migrating or invading through a matrigel matrix for 24h were normalized to untreated cells for comparison with tamoxifen-induced T7 EMT cells (mean +/- SEM, n=3). (**I-L**) Expression levels of epithelial (E-Cadherin, EPCAM) and mesenchymal (MCAM, ECM1) markers in cell-sorted MCF10a-Snail-ER cells. RT-qPCR levels were normalized by TBP expression levels (mean +/- SEM, n=3). *P <0.05, **P <0.01, ***P <0.001 in two-tail paired Student’s t-test respect the corresponding control (empty, dCas9-vSETx2* or negative cells, all in grey).

CRISPR-dCas9 editing systems are known to be heterogenous, with just a percentage of cells properly targeted at the gene locus of interest. To assess the real biological impact of chromatin-induced changes in CTNND1 splicing, we sorted cells in which the mesenchymal-specific protein variant was present, using a splicing-specific antibody recognising only the CTNND1 protein isoform including exon 2 (mCTNND1(ex2)). dCas9-vSETx2-mediated, and thus H3K27me3-driven, induction of exon 2 inclusion in epithelial MCF10a-Snail-ER cells increased the proportion of cells expressing the CTNND1 mesenchymal isoform almost as much as tamoxifen-induced EMT (31% vs 43% positive cells, respectively), whereas the use of the dCas9-vSETx2* mutant had no effect (9% of positive cells as in epithelial cells) (Figure 5G). Moreover, mCTNND1(ex2)-positive cells from dCas9-vSETx2-infected cells, but not dCas9-vSETx2*, included exon 2 at similar levels to tamoxifen-induced EMT cells (Figure S5C), supporting a complete splicing switch to the mesenchymal phenotype only when inducing changes in H3K27 marks. In concordance, EMT was now completely recapitulated, with changes in bi-directional migration, invasion and expression of EMT markers similar to the ones observed in tamoxifen-induced cells (Figure 5H-L). To rule out indirect off-target effects, an independent combination of gRNAs (g2) targeting exon 2 had comparable effects (Figure S5B-G), reinforcing a direct role for chromatin-induced splicing changes in driving important aspects of EMT cell reprogramming.

Collectively, these results support a model by which local changes in exon-specific H3K27 modifications are responsible for the dynamic changes in alternative splicing necessary for cell reprogramming. Moreover, these chromatin-induced changes in splicing are sufficient to induce a change in cell phenotype, providing a novel toolkit for the cell to modulate its proteome in a dynamic and reversible way.

## Discussion

Cell type-specific chromatin and alternative splicing patterns have been intimately involved in cell differentiation and lineage commitment^19, 34^. Increasing evidence has shown a functional cross-talk between these two regulatory layers, whose dysregulation can lead to disease^3, 5, 14, 33^. However, it has remained unclear to what extent chromatin modifications can directly cause cell fate-switching splicing changes. Using CRISPR/dCas9 epigenome editing tools^18^, we have successfully altered local H3K27me3 or H3K27ac levels at alternatively spliced loci. This exon-specific chromatin editing directly affected recognition of these exons by the splicing machinery without affecting overall transcription levels nor RNA Polymerase II kinetics. As we targeted exons essential for the reprogramming of epithelial into mesenchymal cells (EMT), such as CTNND1 exon 2 and FGFR2 exon IIIc, we could show that H3K27-mediated switches in alternative splicing of key EMT exons are sufficient to induce important features of cell reprogramming, demonstrating that chromatin can also regulate cellular identity by driving key changes in alternative splicing patterns.

Changes in splicing-associated histone marks were very dynamic, starting as early as 6h after induction of EMT, even before changes in splicing could be detected. They were also completely reversible, suggesting that epigenetic plasticity could be responsible for the splicing machinery’s dynamic response to a new stimulus, like in EMT. In fact, plants and flies already exploit these chromatin and splicing mediated mechanisms to respond to changes in light and temperature^35–37^. Mammalian cells likely take advantage of the same systems by epigenetically regulating key splicing events for a more efficient and rapid response to external stimuli.

Surprisingly, not all histone marks showed the same dynamics during EMT. For instance, H3K4me1 only correlated with late changes in splicing, similarly to RNA polymerase II elongation rate. Since H3K4me1 levels have been positively associated with RNA polymerase II kinetics^38^, we propose that, contrary to H3K27 marks, H3K4me1 changes could be a consequence of altered splicing, setting up a regulatory feedback loop to reinforce or maintain novel splicing patterns by impacting RNA polymerase II elongation rates.

Chromatin is known to impact splicing by modulating the recruitment of the splicing machinery to weaker RNA binding sites. Several direct physical interactions between splicing and chromatin regulators have been reported. For instance H3K36me3 and H3K9me3 can regulate PTB and SRSF3-dependent splicing via recruitment of the chromatin adaptor proteins MRG15 and HP1, respectively, which by physical interaction favour the binding of the splicing regulators to the pre-mRNA during co-transcriptional splicing^8, 25^. Splicing factors can also interact with the chromatin modifiers, such as hnRNPK with the H3K9 methyltransferase SETDB1 or RBFOX2 with the H3K27 methyltransferase Polycomb Repressor Complex 2^39, 40^. Finally hnRNPA2B1 and hnRNPL were shown to directly interact with chromatin in an RNA-independent way^41^, suggesting that chromatin counts with a variety of molecular mechanisms to impact splicing factors recruitment to the pre-mRNA. Interestingly, histone mark writers, such as p300, have recently been shown to modulate alternative splicing by post-translationally acetylating the splicing factors themselves, which can impact their RNA binding capacity and activity^21^. There is evidence of a p300-mediated acetylation of PTB^42^. However, its functional impact, as well as existence of other post-translational modifications, such as methylation, remain unclear. Since in our particular model system, both acetylation (dCas9-p300, dCas9-Sid4x) and methylation (dCas9-UTX1, dCas9-vSETx2) can affect the same alternatively spliced genes, we consider it unlikely that a splicing factor can be post-translationally regulated by the two marks.

Of note, we do not expect exons marked by a specific histone mark to be dependent on the same splicing regulator, nor all the exons dependent on a specific splicing regulator to be dependent on the same histone mark. In fact, H3K36me3 has been previously shown to modulate recruitment of the splicing repressor PTB and the enhancer SRSF1 at different subsets of alternatively spliced exons^8, 9^. U2 snRNP core splicing regulators can be modulated by H3K4me3 and acetyl H3 marks^10, 11^. Histone acetyltransferases (HATs) and deacetylases (HDACs) have also been shown to differently impact splicing by a variety of mechanisms, from modulating RNA polymerase II elongation rates to directly interacting with splicing regulators such as SF3A1 and SMN1^4, 21, 43^. Finally, of relevance for this work, the aforementioned splicing factor RBFOX2 has recently been shown to induce recruitment of Polycomb Repressive Complex 2 to bivalent gene promoters by protein-protein interactions^40^. Since RBFOX2 and PTB are major splicing regulators of EMT^2^, H3K27me3 enrichment at RBFOX2-dependent exons and H3K27ac enrichment at PTB-dependent sites could represent complementary mechanisms of regulating key splicing events during EMT. In such complex context, confounding proteomics approaches identifying all the protein interactors of a specific histone mark prevalent in the genome, such as H3K27ac, can be limiting to identify novel chromatin/splicing effectors. The development of exon-specific proteomics approaches arises as a promising solution for such mechanistic caveats.

Finally, we expect this chromatin-mediated regulation of alternative splicing to be gene- and context-specific. Recent published genome-wide analysis, in the most extensive epigenomic and transcriptomic datasets publicly available from the ENCODE and Epigenomic Roadmap projects, showed that exons differentially marked by specific histone marks share common functional and regulatory pathways, suggesting a coordinating role for histone marks in regulating the alternative splicing of functionally related exons^12^. In the case of H3K27ac and H3K27me3, we identified 5 genes, intimately involved in cell migration and invasion^17, 33, 44, 45^, which alternative splicing depends on H3K27 marks. Genome-wide studies will be necessary to properly address the global impact of these histone marks in splicing regulation, and to determine what characterizes H3K27-marked exons. Once a list of H3K27ac/me3-dependent exons is identified, we will be better positioned to understand how these histone modifications are regulated during EMT and their impact on cell reprogramming.

In conclusion, we propose that exons sensitive to H3K27 marks might be coregulated during EMT for a rapid induction of changes in splicing necessary for the dynamic functional changes observed during cellular reprogramming. This could have an impact on the development of more specific therapeutic targets to reduce cell invasion and tumour metastasis that depends on EMT phenomena. Therapies targeting general chromatin and splicing factors are currently in use, but often associated with pleiotropic and indirect effects^3, 46^. We propose to use epigenome editing tools to selectively change the splicing-associated chromatin marks responsible for pro-tumorigenic splicing isoforms, such as mCTNND1(ex2). In addition to the H3K27-centric regulation of EMT-related alternative splicing identified here, other histone marks might also coordinate the regulation of alternative splicing events important for other physiological processes. Future studies will bring the necessary insights into this highly dynamic layer of regulation.

## Materials and Methods

### Cell Lines and Cell Culture

#### MCF10a cells

Mcf10a cells are non-transformed human female breast epithelial cells. Mcf10a-Snail-ER cell line was generated by introducing a Snail-1 retroviral expression construct using a fused estrogen receptor (ER) response element to mediate regulation by exogenous 4-hydroxy-tamoxifen (4-OHT) and was obtained from Daniel A. Haber lab with its parental cell line ^1^. All Mcf10a cell lines were maintained at 37°C with 5% CO2 in DMEM/F12 supplemented with 5% horse serum, 10 ng/mL EGF, 10 µg/mL insulin, 0.1 µg/mL cholera toxin, 0.5 µg/mL hydrocortisone, 1% penicillin/streptomycin, 1% L-glutamine (complete medium).

#### EMT induction

Mcf10a were seeded at 7.5.10^5^ cells / 150mm dish and 24h after cells were synchronized in DMEM/F12 supplemented with 10 ng/mL EGF, 10 µg/mL insulin, 0.1 µg/mL cholera toxin, 0.5 µg/mL hydrocortisone, 1% w/v penicillin/streptomycin (No serum medium) for 15h. Cells were then treated with 100nM 4-OHT or Methanol (control) in complete medium.

#### HEK293T cells

HEK293T were maintained at 37°C with 5% CO2 in DMEM supplemented with 10% fetal bovine serum, 1% P/S, 1% L-glutamine. HEK293T are transfected by Calcium Phosphate transfection to generate recombinant lentiviruses.

### Cloning and Plasmids

To generate plasmid DNAs encoding GFP/HAtag epitope-tagged sid4x (a gift from Salton lab), p300core (addgene 61357), EZH2core (a gift from Ducket lab), vSETx2 (from Voigt lab) and UTX core (a gift from Ge lab), the cDNAs were amplified using Q5 High-Fidelity DNA Polymerase (NEB) with primers carrying the appropriate restriction enzymes sites AscI/SbfI (See Table S6 for the list of primers used) and cloned using Quick DNA Ligation Kit (NEB) into dCas9-empty-GFP vector. dCas9-empty-GFP vector has been generated by cutting dCas9-VP64-GFP plasmid (addgene 61422) by BamHI and NheI restriction enzymes to remove VP64 sequence, followed by introduction of a linker containing AscI and SbfI restriction sites and a HAtag epitope-tagged. Q5 Site-Directed Mutagenesis Kit (NEB) was used for generating dCas9 plasmids encoding the mutant p300core* (Y1467F) and vSETx2* (Y105F x2) proteins. Mutagenesis primer sequences and plasmids used in this study are listed in the Table S6. To generate pKLV2.3-Hygro gRNA lentiviral plasmid, the commercial pKLV2.2-PGKpuroBFP plasmid (addgene 72666) was modified by removing the puromycin resistance and the BFP tag, an EcoRI site was added and hygromycin resistance was introduced in XhoI/EcoRI restriction sites. The different gRNAs were cloned by using SapI or BbsI restriction sites. Cloning primer sequences and gRNAs used in this study are listed in the Table S6. Sh RNA plasmids were gifts from different laboratories (See Key Resources Table) or obtained by cloning Sh RNA sequences into pLKO.1-Hygro (addgene 24150) or pLKO.1-Blast (addgene 26655) plasmids with AgeI/EcoRI restriction sites. Sh RNA sequences used in this study are listed in the Table S5.

### Expression and Purification of vSET constructs

The coding sequence fo vSET was ordered from IDT and cloned into a modified pET22b plasmid. Single-chain dimeric vSET constructs with GSGSG-(SSG)n-SGSGG linkers (n=1-3) in between two vSET monomers were generated by PCR and subcloning of a fragment encoding the C-terminal 8 residues of vSET followed by the linker and a complete vSET monomer into the XbaI and HindIII restriction sites of vSET in modified pET22b.

vSET and dimeric sc-vSET (called vSETx2 in this paper) constructs were expressed in BL21 *E. coli* and purified from inclusion bodies essentially as described for vSET by ^47^. In short, inclusion bodies were solubilized in unfolding buffer (20 mM Tris pH 7.5, 7 M guanidine hydrochloride, 10 mM DTT). To refold vSET and vSETx2 proteins, solubilized protein was first dialyzed against urea dialysis buffer (10 mM Tris pH 7.5, 7 M urea, 100 mM NaCl, 1 mM EDTA, 5 mM ß-mercaptoethanol), followed by repeated dilution with vSET refolding buffer (50 mM Tris pH 7.5, 300 mM NaCl, 10% glycerol, 0.1 mM EDTA, 5 mM ß-mercaptoethanol) reducing the concentration of urea from 7 M to 1 M in a step-wise fashion in increments of 1 M (1 h/dialysis step). Finally, refolded vSET and vSETx2 proteins were dialyzed once against vSET refolding buffer and then once against vSET HEPES refolding buffer (50 mM HEPES pH 7.5, 300 mM NaCl, 5% glycerol, 0.1 mM EDTA, 5 mM ß-mercaptoethanol).

Size exclusion chromatography of refolded vSET and vSETx2 constructs was performed on a Superdex 75 column in vSET HEPES refolding buffer.

vSETx2 construct specificity towards H3K27 was tested by methyltransferase assays in which the substrate nucleosomes were harbouring an H3K27A mutation (Figure S2C).

### Methyltransferase assays

In vitro histone methyltransferase (HMT) assays were carried out essentially as described in ^48^. Briefly, core histones were expressed in E. coli, purified from inclusion bodies and assembled into histone octamers by dialysis into refolding buffer (10 mM Tris pH 8, 2 M NaCl, 1 mM EDTA, 5 mM ß-mercaptoethanol). Correctly assembled octamers were purified by size exclusion chromatography on a Superdex S200 column. Recombinant nucleosome arrays were reconstituted via salt dialysis assembly of histone octamers onto plasmid DNA containing 12 177-bp repeats of the 601 nucleosome positioning sequence. To determine methylation activity, 2-10 ng of vSET or VSETx2 constructs were incubated with 1 µg of recombinant nucleosome arrays in 50 mM Tris pH 8.5, 5 mM MgCl2, 4 mM DTT, and ^3^H-labeled SAM for 1 h at 30°C. Reactions were stopped by addition of SDS loading buffer. After separation by SDS-PAGE and transfer to PVDF membranes, loading was assessed by Coomassie staining. Activity was detected as incorporation of ^3^H via exposure of Biomax MS film with the help of Biomax Transcreen LE (both Kodak Carestream) intensifying screens.

### Epigenome Editing

Stable cell lines of MCF10a-Snail-ER expressing the different dCas9s were generated. Briefly, cells were infected with recombinant viruses containing dCas9-empty-GFP, dCas9-sid4x-GFP, dCas9-p300core-GFP, dCas9-EZH2core-GFP, dCas9-vSETx2-GFP or dCas9-UTXcore-GFP following Recombinant Lentivirus Production protocol. Infected cells were harvested and GFP-sorted using a BD FACS Melody (BD Biosciences-US). GFP was excited by a 488-nm laser line and its emission was collected through 527/32BD. dCas9 Stable cell lines were then infected with Lentiviruses containing pKLV2.3-Hygro + gRNAs, were split and medium was supplemented with 100µg/mL hygromycin. See Table S4 for the list of gRNAs used.

### Recombinant Lentivirus Production

HEK393T were split at 2.10^6^ cells / 100mm dish (Day 1). Cells were transfected with 1µg psPAX2 plasmid (VSVG env gene), 1µg pMD2.G plasmid (gag, pol, and accessory proteins), 5µg of plasmid of interest (eg. dCas9-empty), 250mM Cacl2, qsp 500µL sterile water. Samples were gently mixed and completed with 2X HEPES Buffered Saline (HBS), Incubated 15min at room temperature. Mixes were dropped on HEK293T and cells were maintained at 37°C with 5% CO2 (Day 2). 15h after transfection medium was replaced by MCF10a complete medium and MCF10a cells were split at 5.10^5^ cells/100mm dish for further infections (Day 3). 48h and 72h after transfection, viruses were collected, filtered through 0.45µm filter, and dropped on MCF10a cells (Days 4 and 5). 72h after, cells were split and medium was supplemented with 15µg/mL blasticidin or 100µg/mL hygromycin.

### Chromatin Immunoprecipitation

We performed ChIP using H3K27me3 antibody (Cell Signaling C36B11), H3K27Ac antibody (abcam 4729), H3K4me1 antibody (abcam 8895), H3K9Ac antibody (abcam 4441), H3K9me2 antibody (abcam 1220), H3 antibody (Diagenode C15200011), HAtag antibody (abcam 9110), total Pol-II antibody (Santa Cruz sc-55492). MCF10a cells (10 million per sample) were fixed in 1% formaldehyde in PBS at room temperature with agitation for 2min (Histone marks), 4min (HAtag), 10min (Total Pol-II), then quenched with 1M glycine for 5 min. Fixed cells were resuspended in 1mL cold Lysis Buffer A (50mM HEPES pH 7.5, 140mM NaCl, 1mM EDTA, 10% glycerol, 0.5% NP-40/Igepal, 0.25% Triton X-100) prepared fresh with protease inhibitors (Sigma 11836145001) and incubated at 4°C on rotating wheel for 10 min. Nuclei were pelleted and resuspended in 1mL Lysis Buffer B (10mM Tris-HCl pH 8, 200mM NaCl, 1mM EDTA, 0.5mM EGTA, prepared fresh with protease inhibitors), and incubated on rotating wheel for 10 min. Samples were then diluted with 0.75mL Dilution Buffer C (10mM Tris-HCl pH 8, 100mM NaCl, 1mM EDTA, 0.5mM EGTA, 0.1% sodium deoxycholate, 0.5% N-lauroylsarcosine, prepared fresh with protease inhibitors), and sonicated at 4°C for 12, 14, 16 min (for 2, 4, 10 min cross-linking respectively) to generate fragments from 200bp to 1kp long. After sonication, samples were spun at 20,000xg for 30min at 4°C to remove debris. 8µg (Histone marks, HAtag) or 25µg (Total Pol-II) of chromatin were diluted in TSE 150 Buffer (0.1% SDS, 1% Triton X-100, 2mM EDTA, 20mM Tris-HCl pH 8, 150mM NaCl, supplemented with protease inhibitors) and cleaned-up with 30µL of pre-washed Dynabeads Protein G (Thermo Fisher 10009D) and incubated at 4°C on rotating wheel for 1h30. Prior to setting up immunoprecipitation (“IP”) reactions, 50µl of precleared chromatin was removed as “Input.” 150µL of TE/1% SDS Buffer (10mM Tris-HCl pH 8, 1mM EDTA pH 8, 1% SDS) was added to “Input” and incubated overnight at 65°C. 3µl of proteinase K (Thermo Fisher EO0491) was added and samples were incubated at 37°C for 2h. Following the incubation, “Input” DNA was purified using the QIAquick PCR Purification kit (QIAGEN 28106) per the manufacturer’s instructions. To set up IP reactions, precleared chromatin was mixed with antibody and rotated overnight at 4C. IP reactions was added to 30µL pre-washed Dynabeads Protein G and rotated 1h30 at 4°C. Beads were washed once with TSE 150 Buffer, once with TSE 500 Buffer (0.1% SDS, 1% Triton X-100, 2mM EDTA, 20mM Tris-HCl pH 8, 500mM NaCl), once with Washing Buffer (10 mM Tris-HCl pH 8, 1mM EDTA, 0.25 M LiCl, 0.5% NP-40/Igepal, 0.5% sodium deoxycholate), and twice with TE (10mM Tris-HCl pH 8, 1mM EDTA pH 8). Following the final wash, beads were eluted with 100µl of Elution Buffer (50mM Tris-HCl pH 8, 10mM EDTA, 1% SDS) 15min at 65°C while vigorously shaking, and 100µL of TE/1% SDS Buffer for a final eluate volume of 200ul. The following were incubated overnight at 65°C. 3µl of proteinase K was added and samples were incubated at 37°C for 2h. Following the incubation, DNA was purified using the QIAquick PCR Purification kit (QIAGEN 28106) per the manufacturer’s instructions. Input and immunoprecipitated DNA were then analyzed by QPCR using the iTaq Universal Syber Green supermix (Bio-Rad #1725121) on the Bio-Rad CFX-96 Touch Real-Time PCR System. Results are represented as the mean value +/- S.E.M of at least 3 independent experiments of immunoprecipitated chromatin (calculated as a percentage of the input) with the indicated antibodies after normalization by the mean of two control regions stably enriched across the different conditions. See Table S1 for the list of gene-specific primers used.

### RNA Extraction and RT-qPCR

Quantitative RT-PCR analysis was performed in biological triplicates or quadruplicates. Total RNAs were prepared from cells with the GeneJET RNA Purification Kit (Thermo Scientific #K0732). All samples were eluted into 30µl RNAse-free water. DNAs was remove from RNA by using RQ1 RNase-Free DNase (Promega #M6101), briefly, 1µg of RNA was mixed with RQ1 DNase and RQ1 DNase 10X Reaction Buffer and incubated 30min at 37°C. RQ1 enzyme was inactivated by adding Stop Solution 10min at 65°C. cDNAs were generated using the Transcriptor First Strand cDNA Synthesis kit (Roche 04 897 030 001) according to the manufacturer’s instructions. For each biological replicate, quantitative PCR reactions were performed in technical duplicates using the iTaq Universal Syber Green supermix (Bio-Rad) on the Bio-Rad CFX-96 Touch Real-Time PCR System, and the data normalized to *TBP*. Data from biological replicates are plotted as mean +/- S.E.M. See Table S2 for the list of gene-specific primers used.

### Migration and Invasion Assays

#### Nondirectional migration - Wound Healing Assay

MCF10a cells were plated at 1.10^6^ cells/well in 6 well plates containing complete medium and they were grown to confluency. Confluent cultures were serum-starved for 12 hours. Serum-starved, confluent cell monolayers were wounded with a plastic pipette tip and they were washed three times with PBS to remove floating cells. Following washing, the cells were cultured in complete medium. The wounded area was photographed at 0h (control) and 24 hours later using a Zeiss axiovert 40 CFL microscope with a 10X objective (100X magnification). Cell migration into the scratch was quantified using ImageJ plugin MRI Wound Healing Tool (Volker Baecker, Montpellier RIO Imaging).

#### Directional migration – Transwell Filter Assay

Cell migration assay was performed using 24 well chambers (Sigma CLS3422-48EA) with uncoated polycarbonate membranes (pore size 8µm). Briefly, 5.10^4^ cells resuspended in depleted medium (DMEM/F12 supplemented with 1% horse serum, 10 µg/mL insulin, 0.1 µg/mL cholera toxin, 0.5 µg/mL hydrocortisone, 1% penicillin/streptomycin, 1% L-glutamine) were placed in the upper chamber of the transwell unit. The bottom chamber was filled with 0.6mL complete medium supplemented with 20ng/mL FGF-2. The plates were incubated for 12h at 37°C with 5% CO2 and the cells migrating form the upper to the lower chamber of the unit were fixed with 4% paraformaldehyde in PBS 2min, permeabilized with 0.1% Triton X-100 for 5min and stained with 0.2% crystal violet for 1h. Migrating cells were counted using a Zeiss axiovert 40 CFL microscope with a 5X objective (50X magnification).

#### Invasion – Transwell Filter Assay

For cell invasion assay, 24 well chambers were coated with Matrigel (Sigma E6909) diluted in depleted medium for 1h at 37°C and assays were performed as described in Directional migration except the assay was performed for 24h.

### Flow Cytometry Experiments

MCF10a cells were fixed in 4% Paraformaldehyde for 10min at room temperature followed by a 15min permeabilization step in 0.5% Tween20. Cells were resuspended in Blocking Buffer (PBS, 3% BSA, 0.1% Tween20) for 30min on rotating wheel at room temperature and incubated with conjugated antibodies EPCAM-PE (MACS Miltenyi 130-113-264) and MCAM-APC (MACS Miltenyi 130-120-771) for 1h30 on rotating wheel at room temperature protected from light. Cells were harvested and analyzed using a MACS Quant 10 (MACS Miltenyi Biotec). PE was excited by a 488-nm laser line (laser DPSS) and its emission was collected through 655/605nm; APC was excited by a 640-nm laser line and its emission was collected through 655/730nm. The data were analyzed using Flowing software (Perttu Terho, Turku Centre for Biotechnology).

### FACS of CTNND1 Exon 2 Expressing Cells

Cells were resuspended in Blocking Buffer (PBS, 3% BSA) for 30min on rotating wheel at room temperature and successively incubated with CTNND1 e2 primary antibody (Santa Cruz sc-23873) for 1h30 on rotating wheel at room temperature, and PE-Cy7 secondary antibody (Thermo Fisher 25-4015-82) for 30min on rotating wheel at room temperature protected from light. Cells were harvested and analyzed using a BD FACS Melody (BD Biosciences-US). PE-Cy7 was excited by a 561-nm laser line and its emission was collected through 783/56BD. The data were analyzed using BD FACS Chorus software (BD Biosciences-US).

### Polymerase II Elongation Measurement

A DRB treatment (Sigma D1916) of 100µM for 6h was necessary in order to fully block endogenous CFTR transcription. Cells were washed and the kinetic (0, 5, 10, 15, 20, 30, 45, 60, 90 min) was started by adding complete medium. For each time point of the kinetic, cells are scraped and cell pellets are snap frozen in liquid nitrogen. Total RNA was extracted as mentioned above in RNA Extraction and RT-qPCR. Reverse transcriptase reaction was initiated with random hexamers. Quantification of the pre-mRNAs was performed by real-time PCR with amplicons spanning the intron-exon junctions. For each biological replicate, quantitative PCR reactions were performed in technical duplicates using the iTaq Universal Syber Green supermix (Bio-Rad) on the Bio-Rad CFX-96 Touch Real-Time PCR System, and the data normalized by *tRNA*. Data from biological replicates are plotted as mean +/- S.E.M. See Table S3 for the list of gene-specific primers used.

### TSA, Panobinostat and DRB Treatments

A 24 hours treatment of 40µM of DRB (Sigma D1916) or 1µg/mL of TSA (Trichostatin A - Sigma T8552) was applied on MCF10a-Snail-ER cells after 0 days (T0) or 7 days (T7) of EMT induction, to impede the dynamics of transcribing RNA Polymerase II. Total RNA extraction and quantification were performed as mentioned above in Polymerase II Elongation measurement.

For HDAC inhibition during EMT reprogramming, MCF10a-Snail-ER cells were treated with 3µg/mL of TSA (Trichostatin A - Sigma T8552) or 10nM of Panobinostat (gift from Moreaux Lab, IGH) at the same time as addition of Tamoxifen for EMT induction during 24h.

### shRNA Knockdown

Knock-down of HDAC1, HDAC2, PTB, ELAV1, ESRP1, MBNL1, hnRNPFH1, CELF1, hnRNPF, SRSF1, FUS, RBFOX2, SOX9, SMAD3 and PCBP1 was performed according to the Recombinant Lentivirus Production protocol. Briefly, HEK293T cells were transfected with the appropriate shRNA plasmid, 15h after transfection medium was replaced by MCF10a complete medium and MCF10a cells were split for further infections. 48h and 72h after transfection, viruses were collected, filtered through 0.45µm filter, and dropped on MCF10a cells. 72h after, cells were split and medium was supplemented with 15µg/mL blasticidin or 100µg/mL hygromycin.

In the case of the double HDAC1+2 knock-down, cells were infected first with the shRNA against HDAC1, selected using blasticidin, and then infected with a second virus containing the shRNA against HDAC2. After 72h of hygromycin selection, double infected cells were EMT induced with tamoxifen for 24h.

### UV cross-linked RNA-Immunoprecipitation

The day before collection, 10^7^ cells were seeded per condition and IP reaction (5×10^6^ for PTB and 5×10^6^ for normal mouse IgG1 IP’s) in a p15 plate. Next, day, cell media was discarded and each plate was washed with 12 ml of cold PBS 1X (D8537, Sigma-Aldrich). Cells were UV-crosslinked at 254 nm with 2000 J/m^2^ in ice and scrapped. After centrifugation at 2500 rpm for 5 min, the PBS was discarded and the pellets were stored at −80 °C until processing. Cells were lysed in 617.5 µl of cell lysis buffer (1% v/v NP-40, 400 U/ml of RNAse inhibitor in 1x PBS) for 10 min in ice. Sodium deoxycholate was added to 0.5% v/v final concentration and samples were incubated with rotation for 15 min at 4°C. Samples were incubated at 37 °C with 30 U of DNAse with shacking at 300 rpm, vortexed briefly and sonicated for 10 cycles x (30’’ on /30’’ off, high setting condition) in 15 ml conical polystyrene tubes using a Bioruptor ^TM^ (Diagenode) sonicator with a 4°C water bath cold circulation system. After that, the tubes were spun to recover all sample and centrifuged for 15 min at 21,130 rcf to remove insoluble debris. Every sample was divided in 2 x 300 µl aliquots and 30 µl were saved as « Input » control and stored at −80 °C. Every aliquot was incubated with 6µg of a-PTB (Ref. 32-4800, Invitrogen) or 6µg of a-normal mouse control IgG1 (Ref. 14-4714-82, Invitrogen), o/n with rotation at 4°C. Next day, 40 µl of Dynabeads protein G (Ref. 10009D, Invitrogen), pre-washed three times with 1ml 1xPBS 0.01% v/v Tween-20, were added per sample and incubated for 4h at 4°C with rotation. The unbound supernatant was discarded and beads were washed once with Cell Lysis Buffer (1% v/v NP-40, 0.5% sodium deoxycholate in 1X PBS), three times with Washing Buffer I (1% v/v NP-40, 0.5% sodium deoxycholate, 300mM NaCl in 1X PBS), once with Washing Buffer II (0.5% v/v NP-40, 0.5% sodium deoxycholate, 0.125% v/v SDS in 1X PBS) and once in PBS 1X. All washes were done for 5 min with rotation at 4°C. Beads and Inputs were incubated with 100µl of Proteinase K buffer (100 mM Tris-HCl pH 8, 50 mM NaCl, 10 mM EDTA, 0.5% v/v SDS, 100 U RNAsin of in DEPC water) and 10 µl of Proteinase K for 45 min at 25°C shacking at 1,500 rpm. 1ml of Trizol® (Ambion) was added per sample (beads or input) and RNA was purified according to manufacturer’s protocol, including 1µl of Glycoblue^TM^ Coprecipitant (Ref. AM9515, Invitrogen). RNA pellets were resuspended in 8µl of DEPC water and incubated with µl (1U DNAse) and 1 µl of 10X DNAse buffer (Ref. M6101, Promega) for 30 min at 37°C. The DNAse was inactivated with 1 µl of Stop Solution at 65°C for 10 min and RT was performed using Transcriptor First Strand cDNA Synthesis Kit (Ref. 04 897 030 001, Roche) in a final volume of 20 µl. RT was diluted 1/5 and each sample was quantified in duplicates as described before. The enrichment of every IP was normalized to its Input using (2^(Ct IP-Ct Input) and for representation the fold change was calculated relative to IgG enrichment.

### Motif search analysis

RNA binding motif search analysis was done using CTNND1 exon2 sequence in four public softwares: RBPDB v1.3 (http://rbpdb.ccbr.utoronto.ca), RBPMAP v1.1 (http://rbpmap.technion.ac.il), SFMAP v1.8 (http://sfmap.technion.ac.il/), Spliceaid (http://www.introni.it/splicing.html). All softwares were used with the default parameter settings, except for some exceptions. For RBPDB the threshold 0.8 was applied. For RBPMAP, the Stringency level used was “High stringency” with all motifs available from Human/mouse. For SFMAP both “Perfect match” and “High stringency” stringency levels were used. For catRAPID, the following settings were used: “Full-length proteins”, “RNA and DNA binding” and including “disordered proteins. We prioritized the RNA motifs predicted by more than one database and expressed in MCF10a cells.

### P Values and Statistical Analysis

Two-tailed paired Student’s t-test was used in all Figures and Supplementary Figures. P-values and other details can be found in figure legends.

## Acknowledgments

We are thankful to Dr. Haber for the EMT cellular model, Dr. Bertrand for the modified gRNA plasmid, Dr. Salton for plasmid reagents, Dr. Duckett for EZH2 plasmid, Dr. Pradeepa for Sid4x plasmid and Dr. Ge for UTX1 plasmid. We are also thankful to Paola Scaffidi, Andrew Oldfield, and Bernard de Massy for critical reading and discussion of the manuscript.

## Author contributions

Conceptualization: A.S., Y.N.A. and R.L.; Methodology & Investigation: A.S., Y.N.A. and K.W.; Resources: P.V.; Writing & editing: A.S., Y.N.A., A.O., P.V. and R.L.; Funding Acquisition: P.V. and R.L.

## Data and materials availability

see Resources Table

## Resources table

Supplementary List: List of Reagents and ressources

Supplementary Table S1: List of ChIP-qPCR primers

Supplementary Table S2: List of RT-qPCR primers

Supplementary Table S3: List of Polymerase II elongation assay and RNA-IP primers

Supplementary Table S4: List of gRNAs

Supplementary Table S5: List of shRNAs

Supplementary Table S6: List of cloning primers

---

**Supplementary Figure 1:**
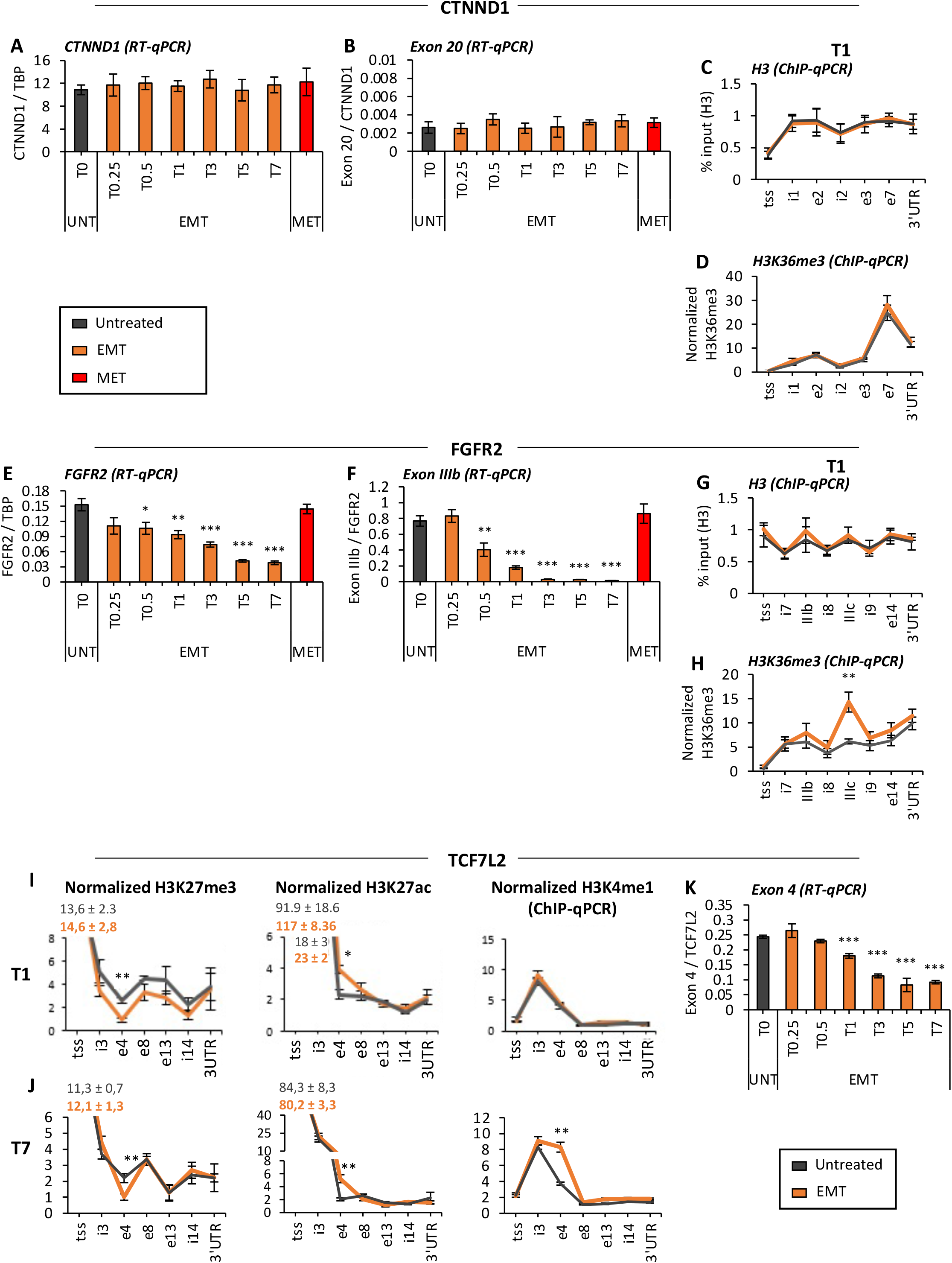
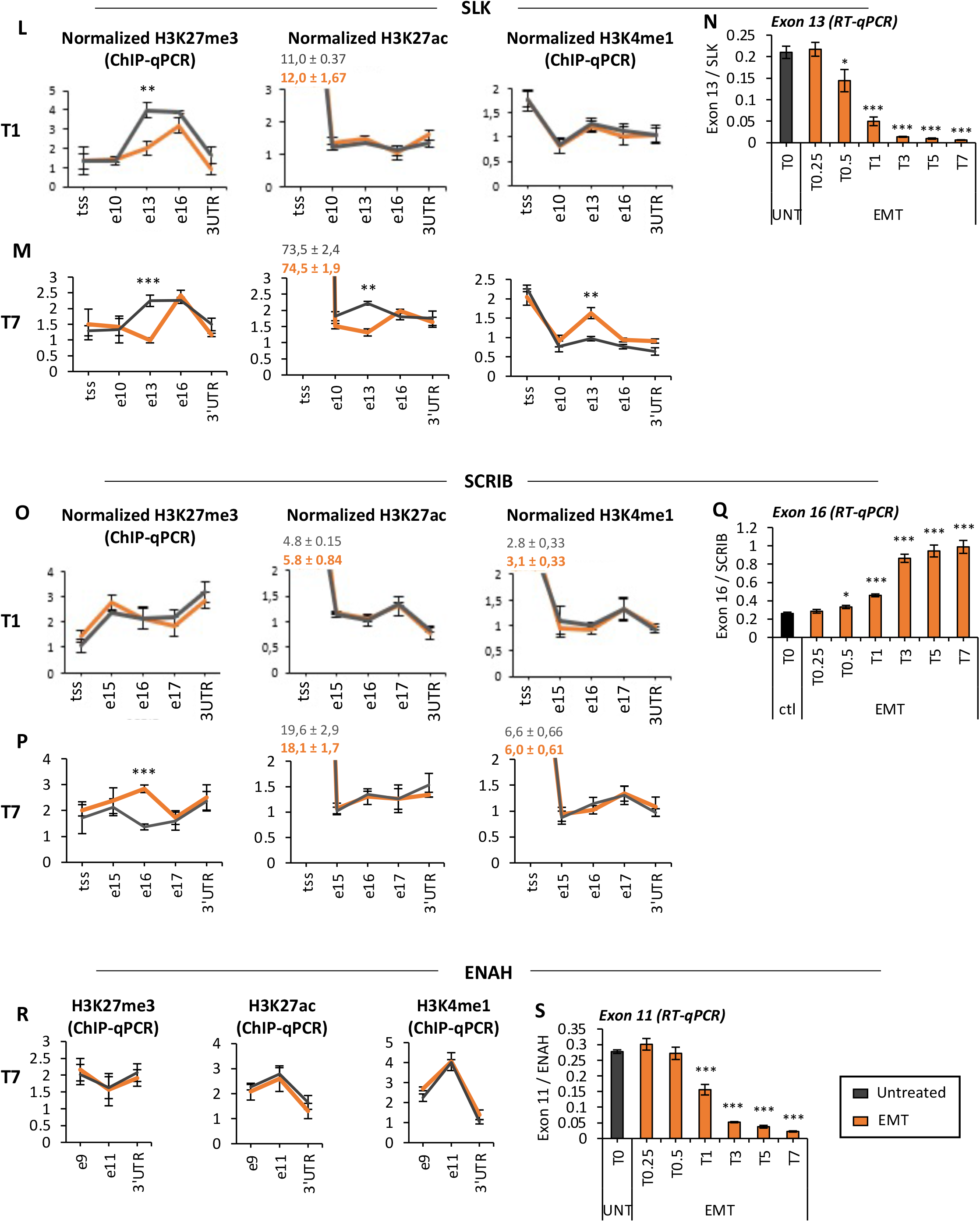
Localised enrichment of specific histone marks at alternatively spliced exons during EMT. (**A,B**) CTNND1 expression and exon 20 inclusion levels relative to total TBP and CTNND1, respectively, during EMT and MET in induced MCF10a-Snail-ER cells by RT-qPCR (mean +/- SEM, n=4). (**C,D**) Enrichment levels of H3 and H3K36me3 along CTNND1 locus in tamoxifen-induced MCF10a-Snail-ER cells for 24h by ChIP-qPCR (mean +/- SEM, n=4). Only for H3K36me3, the percentage of input was normalized by two control regions across the different conditions. (**E,F**) FGFR2 expression and exon IIIb inclusion levels relative to total TBP and FGFR2, respectively, during EMT and MET in induced MCF10a-Snail-ER cells by RT-qPCR (mean +/- SEM, n=4). (**G,H**) Enrichment levels of total H3 and H3K36me3 along FGFR2 locus in tamoxifen-induced MCF10a-Snail-ER cells for 24h by ChIP-qPCR (mean +/- SEM, n=4). Only for H3K36me3, the percentage of input was normalized by two control regions across the different conditions (**I,J,L,M,O,P,R**) Enrichment levels of H3K27me3, H3K27ac and H3K4me1 along TCF7L2 (I-J), SLK (L-M), SCRIB (O-P) and ENAH (R) loci in tamoxifen-induced MCF10a-Snail-ER cells for 1 (T1) or 7 (T7) days by ChIP-qPCR (mean +/- SEM, n=4). The percentage of input was normalized by two control regions across the different conditions. (**K,N,Q,S**) Inclusion levels of alternatively spliced exons essential for EMT: TCF7L2 exon 4 (K), SLK exon 13 (N), SCRIB exon 16 (Q) and ENAH exon 11 (S) in MCF10a-Snail-ER during induction of EMT and reversible MET. RT-qPCR values were normalized by total expression levels of SLK, TCF7L2, SCRIB and ENAH, respectively (mean +/- SEM, n=4). *P <0.05, **P <0.01, ***P <0.001 in two-tail paired Student’s t-test respect untreated (grey).

**Supplementary Figure 2:**
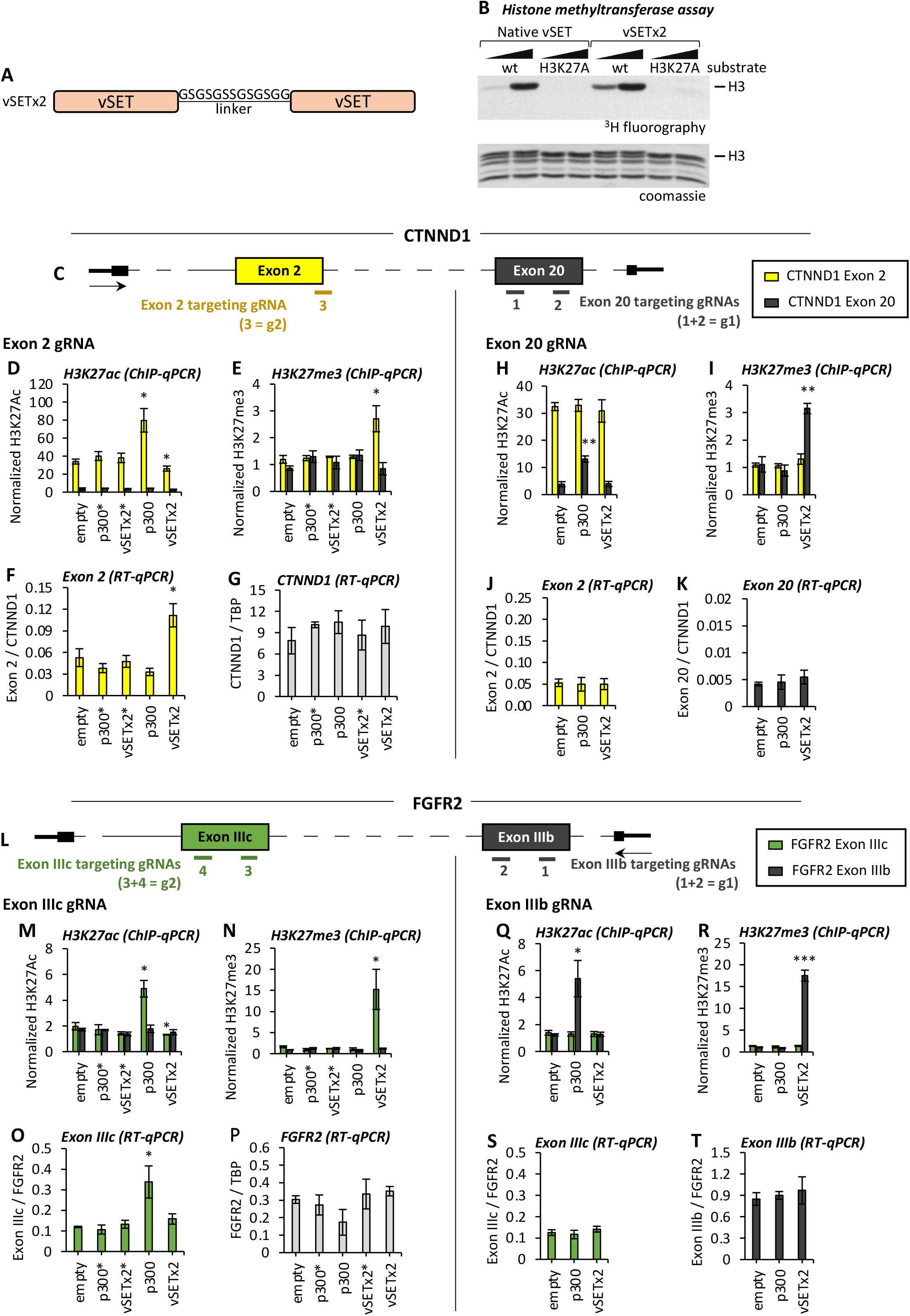
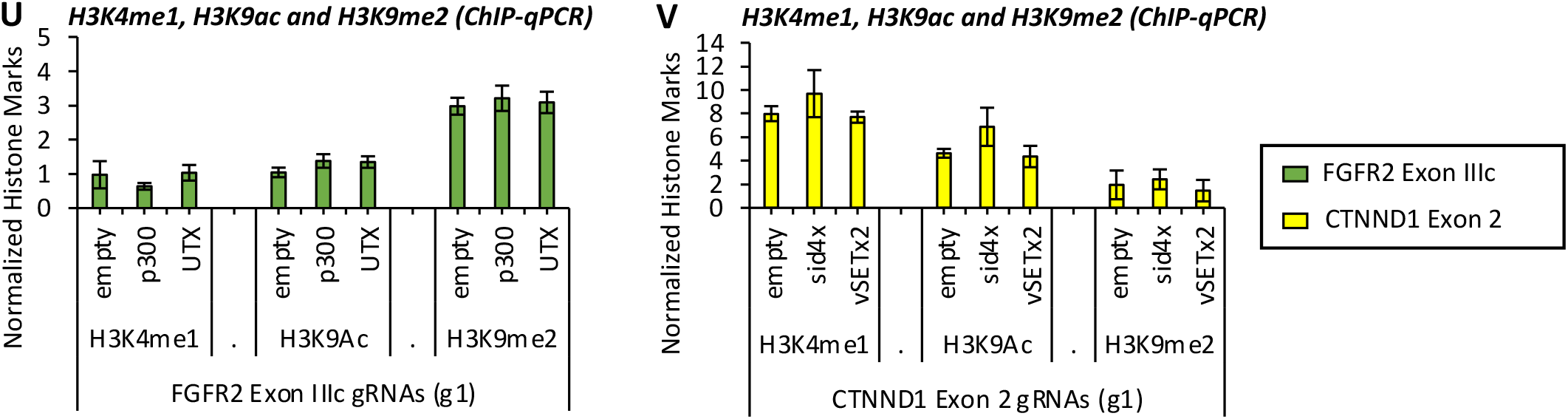
Exon-specific epigenome editing of H3K27 marks is sufficient to induce a change in splicing. (**A**) Schematic of vSETx2 construct with optimal linker sequence between the two monomers. (**B**) Representative histone methyltransferase assay to show the specificity of native vSET and vSET2x activity on H3K27 residue in wild-type and H3K27A mutated recombinant chromatin templates. (**C**) Schematic representation of CTNND1 gene locus and the alternatively spliced exons 2 (yellow) and control exon 20 (grey) with the position of the gRNAs used to exon-specifically target the different dCas9-fused proteins. (**D,E,H,I**) Enrichment levels of H3K27ac (D,H) and H3K27me3 (E,I) at CTNND1 exon 2 (yellow) and control exon 20 (grey) in MCF10a-Snail-ER cells infected with dCas9-fused proteins and two different combination of exon-specific gRNAs targeting exon 2 (g2) or exon 20 (g1) (mean +/- SEM, n=3). The percentage of input was normalized by two control regions across the different conditions. (**F,G,J,K**) Expression levels of CTNND1 exon 2 (F,J), total CTNND1 (G) and exon 20 (K) in untreated MCF10a-Snail-ER cells infected with dCas9-fused proteins and the exon-specific gRNAs targeting exon 2 (g2) or exon 20 (g1). RT-qPCR values were normalized by total CTNND1 or TBP as indicated in the graph (mean +/- SEM, n=4). (**L-T**) Same as (C-K) on FGFR2 gene locus, with (**L**) a schematic representation of FGFR2 locus, with gRNAs position at the alternatively spliced exon IIIc (green) and control IIIb (grey). (**N-T**) H3K27ac, H3K27me3 and expression levels as represented in (D-K) (mean +/- SEM, n=4). (**U,V**) Enrichment levels of H3K4me1, H3K9ac and H3K9me2 at FGFR2 exon IIIc (U, green) or CTNND1 exon 2 (V, yellow) in untreated MCF10a-Snail-ER cells infected with dCas9-fused proteins and exon-specific gRNAs targeting exon IIIc (g1,U) or exon 2 (g1,V) by ChIP-qPCR (mean +/- SEM, n=3). The percentage of input was normalized by two control regions across the different conditions. *P <0.05, **P <0.01, ***P <0.001 in two-tail paired Student’s t-test compared to empty-dCas9.

**Supplementary Figure 3:**
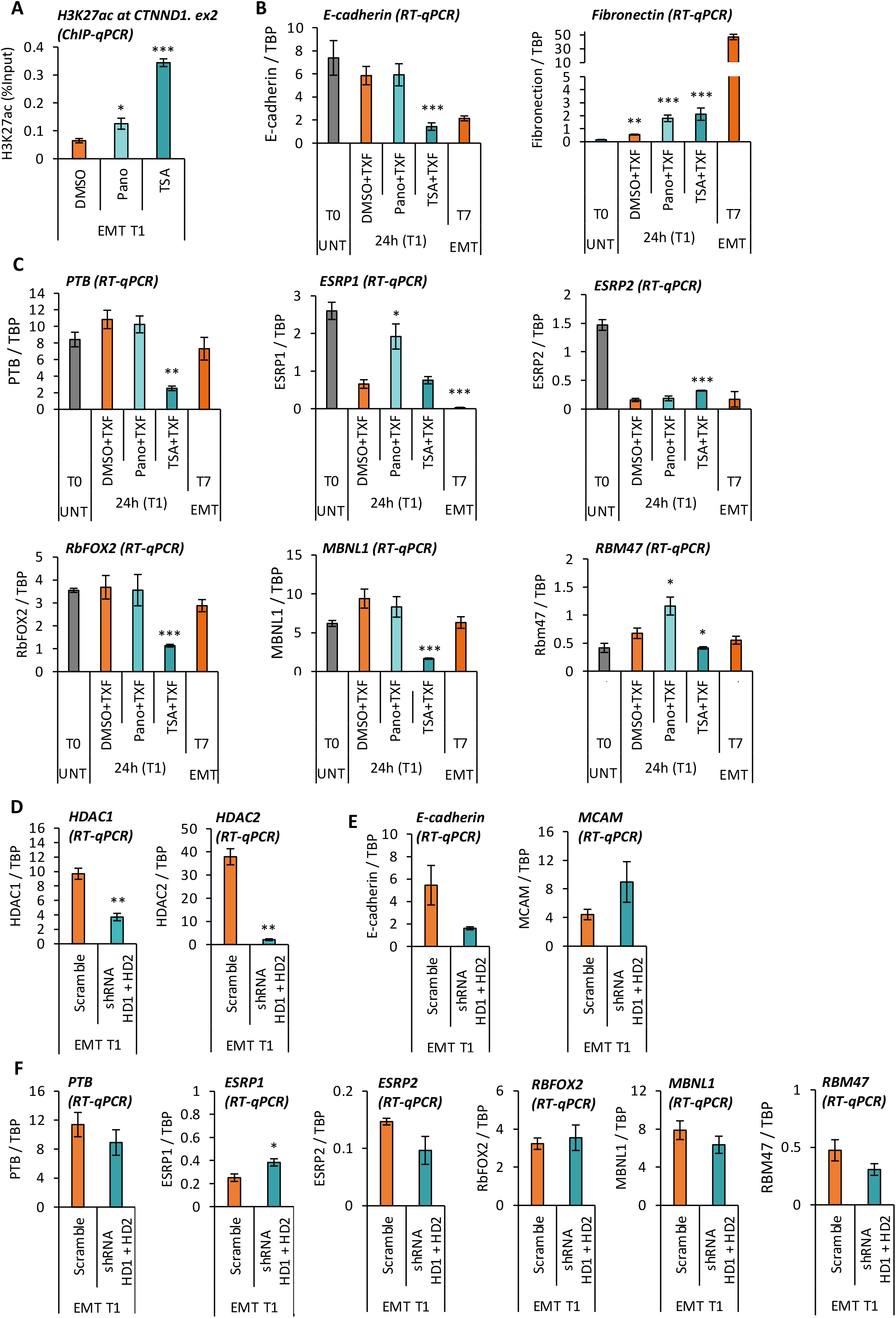
Impact of HDAC inhibition in H3K27 marks and major splicing regulators. (**A**) Enrichment levels of H3K27ac at CTNND1 exon 2 in MCF10a-Snail-ER cells treated for 24h with vehicle DMSO, 10nM of Pano (Panobinostat) or 1µg/mL of TSA (Trichostatin A) during EMT induction with tamoxifen. ChIP-qPCR data is shown as the percentage of input (mean +/- SEM, n=3). (**B-C**) Expression levels of epithelial (E-cadherin) and mesenchymal (Fibronectin) markers (B) and key EMT splicing factors (C) in MCF10a-Snail-ER cells treated for 24h with vehicle DMSO, 10nM of Pano (Panobinostat) or 1µg/mL of TSA (Trichostatin A) during EMT induction with tamoxifen. Untreated (T0) and fully induced EMT cell (T7) are shown as control references. RT-qPCR data was normalized by TBP (mean +/- SEM, n=3). **(D, E, F)** Expression levels of HDAC1, HDAC2 (D), EMT markers (E) and splicing factors (F) upon double knock-down of HDAC1 (HD1) and HDAC2 (HD2) in MCF10a-Snail-ER cells induced for EMT during 24h. Non-targeting scramble shRNA was used as a control. RT-qPCR data was normalized by TBP (mean +/- SEM, n=3). *P <0.05, **P <0.01, ***P <0.001 in two-tail paired Student’s t-test respect T1 DMSO for (A-D) and T1 Scramble for (E-G).

**Supplementary Figure 4.**
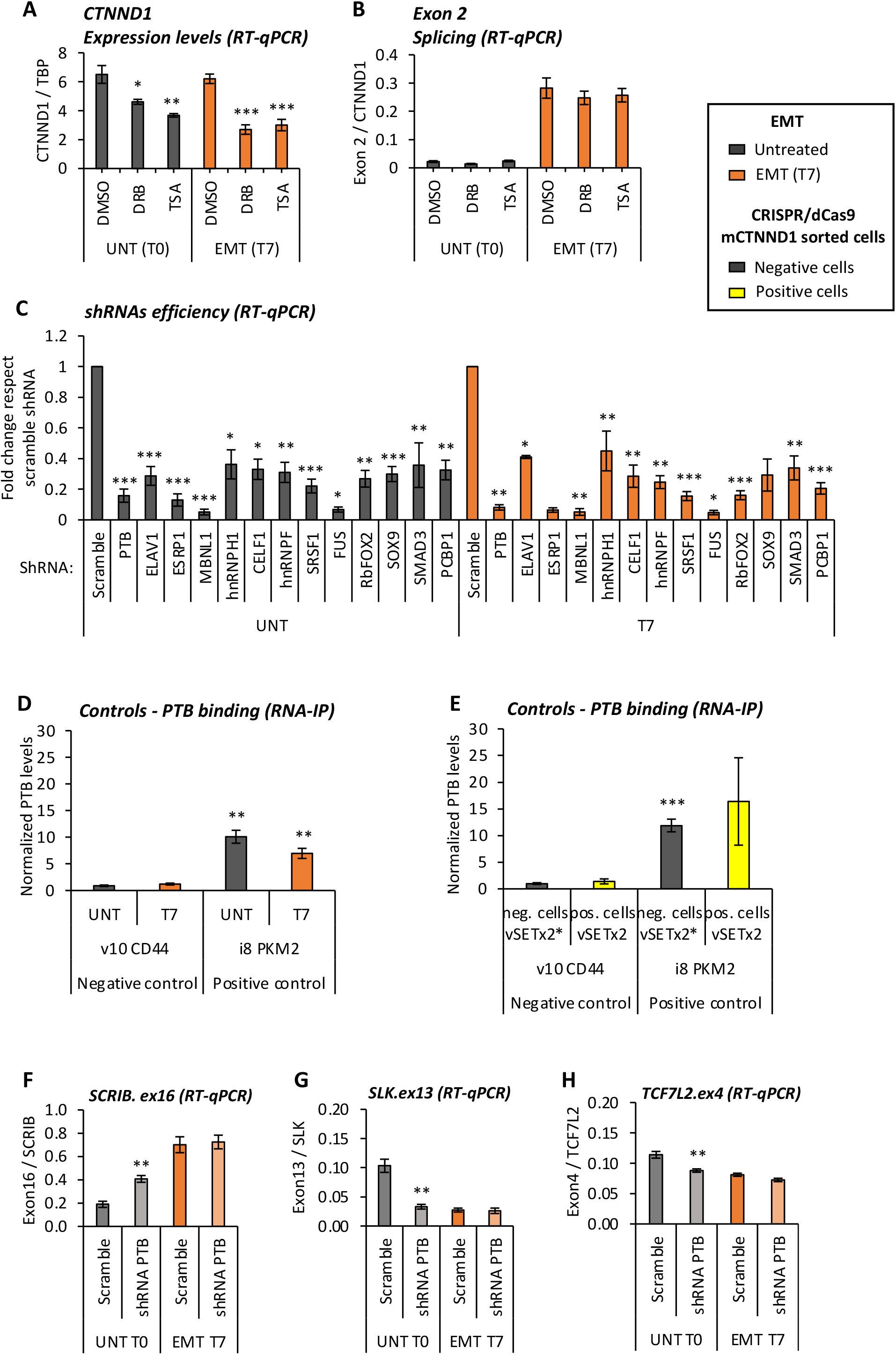
H3K27 marks regulate splicing by modulating the recruitment of RNA-binding proteins, such as PTB. (**A,B**) CTNND1 expression and exon 2 inclusion levels normalized by total TBP or CTNND1 expression levels, respectively, in untreated epithelial (UNT, grey) and mesenchymal-like (EMT T7, orange) MCF10a-Snail-ER cells treated with DMSO (control), 1 µg/mL TSA (HDAC inhibitor) or 40µM DRB (RNA Polymerase II inhibitor) for 24h. RT-qPCR results are shown as the mean +/- SEM of n=4 biological replicates. (**C**) Total expression levels of the candidate splicing factors involved in CTNND1 exon 2 regulation upon shRNA knockdown in untreated (UNT, grey) and tamoxifen-induced (T7, orange) MCF10a-Snail-ER cells. RT-qPCR levels are shown relative to cells infected with scramble shRNA (mean +/- SEM, n=3). **(D)** PTB enrichment levels at the the negative control CD44 v10 and positive control PKM2 intron 8 in untreated (UNT) and tamoxifen-induced (T7) MCF10a-Snail-ER cells. The percentage of input of UV-crosslinking RNA immunoprecipitations were normalized by IgG and CTNND1 exon 7 control levels as in Figure 4 (mean ± SEM, n=5). (**E**) PTB enrichment levels at the same positive and negative control regions as in (D) in cell-sorted MCF10-Snail-ER cells expressing (positive) or not (negative) the mesenchymal-specific splicing isoform mCTNND1(ex2) upon infection with dCas9-vSETx2, or mutant dCas9-vSETx2*, and the exon-specific gRNAs (g1) targeting CTNND1 exon 2. The percentage of input of UV-crosslinking RNA immunoprecipitations were normalized by IgG and CTNND1 exon 7 control levels (mean ± SEM, n=4). *P <0.05, **P <0.01 in two-tail paired Student’s t-test respect the negative control CD44 v10. (**F-H**) Inclusion levels of the indicated exons relative to the total expression levels of their corresponding gene upon shRNA knock-down of PTB in epithelial untreated (UNT T0, grey) and tamoxifen-induced (EMT T7, orange) MCF10a-Snail-ER cells. Non-targeting shRNA is used as a control (Scramble)(mean +/- SEM, n=4). *P <0.05, **P <0.01, ***P <0.001 in two-tail paired Student’s t-test respect cells infected with scramble shRNA.

**Supplementary Figure 5:**
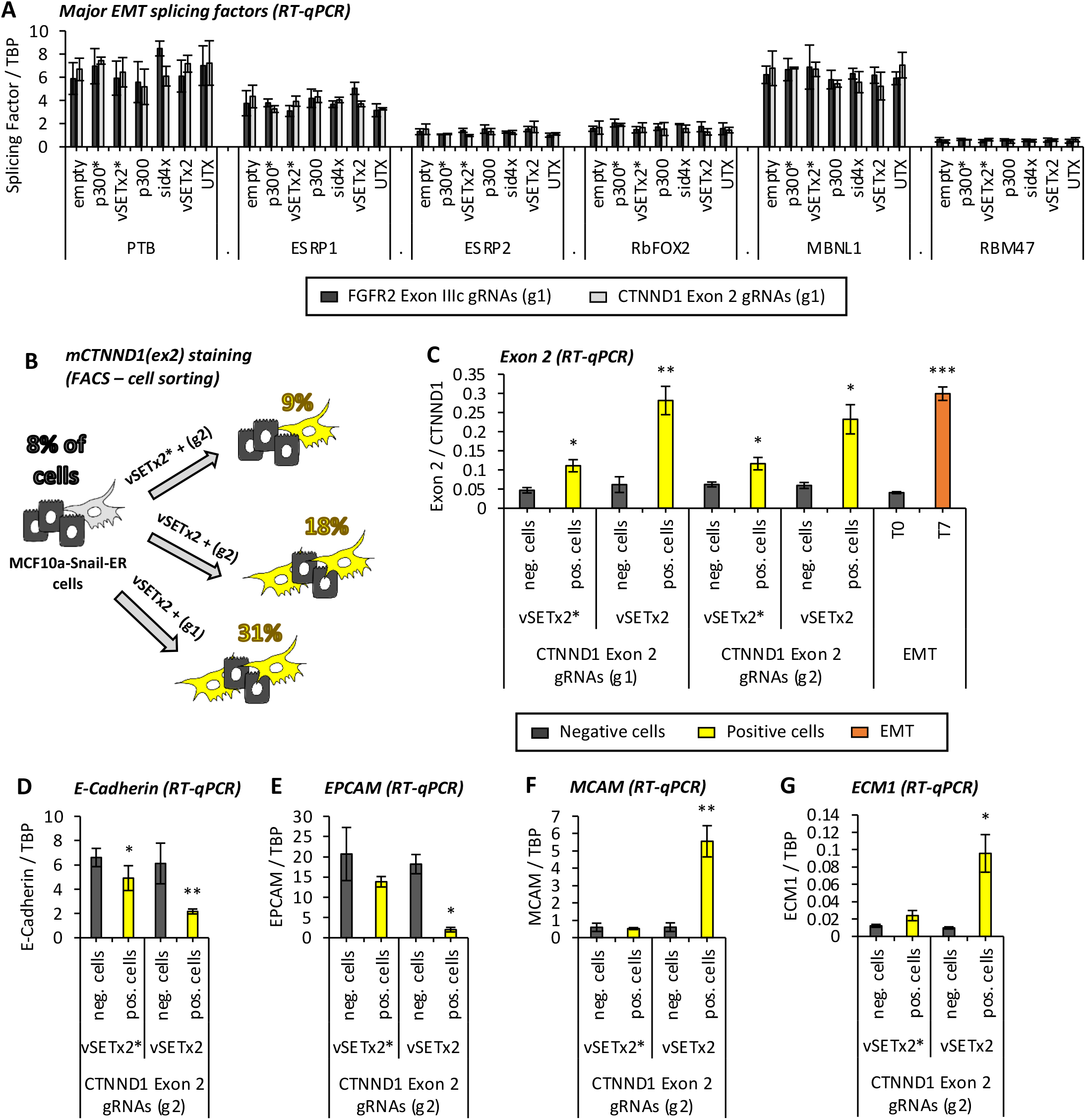
Direct effect of dCas9 epigenomic editing on EMT. (**A**) Expression levels of the splicing factors most important for EMT (PTB, ESRP1, ESRP2, RbFOX2, MBNL1 and RBM47) in MCF10a-Snail-ER cells infected with different dCas9-fused proteins and exon-specific gRNAs targeting FGFR2 exon IIIc (g1, dark grey) or CTNND1 exon 2 (g1, light grey). RT-qPCR levels were normalized by TBP (mean +/- SEM, n=3). (**B**) MCF10a-Snail-ER cells infected with dCas9-vSETx2 or mutant dCas9-vSETx2* and two different combinations of exon-specific gRNAs (g1 and g2) targeting CTNND1 exon 2 were cell-sorted by expression levels of the mesenchymal CTNND1 protein variant, which includes exon 2 (mCTNND1(ex2)), using splicing-specific antibodies. Negative cells not expressing mCTNND1(ex2) and tamoxifen-induced T7 EMT cells were used as controls. The percentage of mCTNND1(ex2) positive cells per condition is shown. (**C**) CTNND1 exon 2 inclusion levels in cells expressing (positive) or not (negative) the splicing variant mCTNND1(ex2) in the conditions described in (B). RT-qPCR levels were normalized by total CTNND1 expression levels (mean +/- SEM, n=3). (**D-G**) Expression levels of epithelial (E-Cadherin, EPCAM) and mesenchymal (MCAM, ECM1) markers in cell-sorted MCF10a-Snail-ER cells infected with dCas9-vSETx2 or the mutant dCas9-vSETx2* and the second combination (g2) of gRNAs targeting CTNND1 exon2. RT-qPCR levels were normalized by TBP expression levels (mean +/- SEM, n=3). *P <0.05, **P <0.01, ***P <0.001 in two-tail paired Student’s t-test respect negative cells.

## References and Notes

1. Javaid, S. et al. Dynamic Chromatin Modification Sustains Epithelial-Mesenchymal Transition following Inducible Expression of Snail-1. Cell reports 5, 1679–89 (2013).

2. Shapiro, I. M. et al. An EMT-driven alternative splicing program occurs in human breast cancer and modulates cellular phenotype. PLoS Genet 7, e1002218 (2011).

3. Daguenet, E., Dujardin, G. & Valcarcel, J. The pathogenicity of splicing defects: mechanistic insights into pre-mRNA processing inform novel therapeutic approaches. EMBO Rep 16, 1640–55 (2015).

4. de la Mata, M. et al. A slow RNA polymerase II affects alternative splicing in vivo. Mol Cell 12, 525–32 (2003).

5. Braunschweig, U., Gueroussov, S., Plocik, A. M., Graveley, B. R. & Blencowe, B. J. Dynamic integration of splicing within gene regulatory pathways. Cell 152, 1252–69 (2013).

6. Luco, R. F., Allo, M., Schor, I. E., Kornblihtt, A. R. & Misteli, T. Epigenetics in alternative pre-mRNA splicing. Cell 144, 16–26 (2011).

7. Guo, R. et al. BS69/ZMYND11 reads and connects histone H3.3 lysine 36 trimethylation-decorated chromatin to regulated pre-mRNA processing. Mol Cell 56, 298–310 (2014).

8. Luco, R. F. et al. Regulation of alternative splicing by histone modifications. Science 327, 996–1000 (2010).

9. Pradeepa, M. M., Sutherland, H. G., Ule, J., Grimes, G. R. & Bickmore, W. A. Psip1/Ledgf p52 Binds Methylated Histone H3K36 and Splicing Factors and Contributes to the Regulation of Alternative Splicing. PLoS Genet 8, e1002717 (2012).

10. Sims, R. J. et al. Recognition of trimethylated histone H3 lysine 4 facilitates the recruitment of transcription postinitiation factors and pre-mRNA splicing. Mol Cell 28, 665–76 (2007).

11. Gunderson, F. Q. & Johnson, T. L. Acetylation by the transcriptional coactivator Gcn5 plays a novel role in co-transcriptional spliceosome assembly. PLoS Genet 5, e1000682 (2009).

12. Agirre, E., Oldfield, A., Bellora, N., Segelle, A. & Luco, R. F. Splicing-associated chromatin signatures: a combinatorial and position-dependent role for histone marks in splicing definition. Nature comm 12, 682 (2021).

13. Li, T., Liu, Q., Garza, N., Kornblau, S. & Jin, V. X. Integrative analysis reveals functional and regulatory roles of H3K79me2 in mediating alternative splicing. Genome Med 10, 30 (2018).

14. Xu, Y., Zhao, W., Olson, S. D., Prabhakara, K. S. & Zhou, X. Alternative splicing links histone modifications to stem cell fate decision. Genome Biol 19, 133 (2018).

15. Gonzalez, I. et al. A lncRNA regulates alternative splicing via establishment of a splicing-specific chromatin signature. Nat Struct Mol Biol 22, 370–6 (2015).

16. Ranieri, D. et al. Expression of the FGFR2 mesenchymal splicing variant in epithelial cells drives epithelial-mesenchymal transition. Oncotarget 7, 5440–60 (2016).

17. Yanagisawa, M. et al. A p120 catenin isoform switch affects Rho activity, induces tumor cell invasion, and predicts metastatic disease. J Biol Chem 283, 18344–54 (2008).

18. Hilton, I. B. et al. Epigenome editing by a CRISPR-Cas9-based acetyltransferase activates genes from promoters and enhancers. Nat Biotechnol 33, 510–7 (2015).

19. Margueron, R. & Reinberg, D. The Polycomb complex PRC2 and its mark in life. Nature 469, 343–9 (2011).

20. Hong, S. et al. Identification of JmjC domain-containing UTX and JMJD3 as histone H3 lysine 27 demethylases. PNAS 104, 18439–18444 (2007).

21. Siam, A. et al. Regulation of alternative splicing by p300-mediated acetylation of splicing factors. RNA 25, 813–824 (2019).

22. Mujtaba, S. et al. Epigenetic transcriptional repression of cellular genes by a viral SET protein. Nat. Cell Biol. 10, 1114–1122 (2008).

23. Ha, K. et al. Histone deacetylase inhibitor treatment induces ‘BRCAness’ and synergistic lethality with PARP inhibitor and cisplatin against human triple negative breast cancer cells. Oncotarget 5, 5637–5650 (2014).

24. Montgomery, R. L. et al. Histone deacetylases 1 and 2 redundantly regulate cardiac morphogenesis, growth, and contractility. Genes Dev. 21, 1790–1802 (2007).

25. Yearim, A. et al. HP1 is involved in regulating the global impact of DNA methylation on alternative splicing. Cell reports 10, 1122–34 (2015).

26. Young, J. I. et al. Regulation of RNA splicing by the methylation-dependent transcriptional repressor methyl-CpG binding protein 2. Proc Natl Acad Sci U S A 102, 17551–8 (2005).

27. Girardot, M. et al. SOX9 has distinct regulatory roles in alternative splicing and transcription. Nucleic Acids Res 46, 9106–9118 (2018).

28. Tripathi, V. et al. Direct Regulation of Alternative Splicing by SMAD3 through PCBP1 Is Essential to the Tumor-Promoting Role of TGF-1. Mol. Cell 64, 549–564 (2016).

29. Warzecha, C. C. et al. An ESRP-regulated splicing programme is abrogated during the epithelial-mesenchymal transition. EMBO J 29, 3286–300 (2010).

30. Carstens, R. P., Eaton, J. V., Krigman, H. R., Walther, P. J. & Garcia-Blanco, M. A. Alternative splicing of fibroblast growth factor receptor 2 (FGF-R2) in human prostate cancer. Oncogene 15, 3059–65 (1997).

31. Sebestyén, E., Zawisza, M. & Eyras, E. Detection of recurrent alternative splicing switches in tumor samples reveals novel signatures of cancer. Nucleic Acids Res 43, 1345–1356 (2015).

32. Villemin, J.-P. et al. A cell-to-patient machine learning transfer approach uncovers novel basal-like breast cancer prognostic markers amongst alternative splice variants. BMC Biology 19, 70 (2021).

33. Sanidas, I. et al. Phosphoproteomics screen reveals akt isoform-specific signals linking RNA processing to lung cancer. Mol Cell 53, 577–90 (2014).

34. Gabut, M. et al. An alternative splicing switch regulates embryonic stem cell pluripotency and reprogramming. Cell 147, 132–46 (2011).

35. Martin Anduaga, A. et al. Thermosensitive alternative splicing senses and mediates temperature adaptation in Drosophila. eLife 8, e44642 (2019).

36. Pajoro, A., Severing, E., Angenent, G. C. & Immink, R. G. H. Histone H3 lysine 36 methylation affects temperature-induced alternative splicing and flowering in plants. Genome Biol 18, 102 (2017).

37. Petrillo, E. et al. A chloroplast retrograde signal regulates nuclear alternative splicing. Science 344, 427–430 (2014).

38. Jonkers, I., Kwak, H. & Lis, J. T. Genome-wide dynamics of Pol II elongation and its interplay with promoter proximal pausing, chromatin, and exons. eLife 3, e02407 (2014).

39. Thompson, P. J. et al. hnRNP K coordinates transcriptional silencing by SETDB1 in embryonic stem cells. PLoS Genet 11, e1004933 (2015).

40. Wei, C. et al. RBFox2 Binds Nascent RNA to Globally Regulate Polycomb Complex 2 Targeting in Mammalian Genomes. Mol Cell 62, 875–889 (2016).

41. Kfir, N. et al. SF3B1 association with chromatin determines splicing outcomes. Cell reports 11, 618–29 (2015).

42. Weinert, B. T. et al. Time-Resolved Analysis Reveals Rapid Dynamics and Broad Scope of the CBP/p300 Acetylome. Cell 174, 231–244.e12 (2018).

43. Rahhal, R. & Seto, E. Emerging roles of histone modifications and HDACs in RNA splicing. Nucleic Acids Res 47, 4911–4926 (2019).

44. Roovers, K. et al. The Ste20-like kinase SLK is required for ErbB2-driven breast cancer cell motility. Oncogene 28, 2839–2848 (2009).

45. Wenzel, J. et al. Loss of the nuclear Wnt pathway effector TCF7L2 promotes migration and invasion of human colorectal cancer cells. Oncogene 39, 3893–3909 (2020).

46. Ellis, L., Atadja, P. W. & Johnstone, R. W. Epigenetics in cancer: targeting chromatin modifications. Mol Cancer Ther 8, 1409–20 (2009).

47. Manzur, K. L. et al. A dimeric viral SET domain methyltransferase specific to Lys27 of histone H3. Nat. Struct. Biol. 10, 187–196 (2003).

48. Voigt, P. et al. Asymmetrically modified nucleosomes. Cell 151, 181–193 (2012).

